# Immune Cell Attachment on Material Surface Promotes Bacterial Aggregation and Biofilms

**DOI:** 10.1101/2024.10.28.620772

**Authors:** Rakesh Kumar Pradhan, Sameer Kumar Jagirdar, Kaushika Kodieswaran, Sahana Kumar, Shruthi Ksheera Sagar, Bishal Kumar Nahak, Arshad Khan, Zong-Hong Lin, Balasubramanian Gopal, Siddharth Jhunjhunwala

## Abstract

Bacterial biofilms on indwelling medical devices is a major driver of healthcare-associated infection despite significant advances in antifouling surface engineering, suggesting that laboratory antibacterial performance does not fully capture the determinants of clinical colonization. Here we show that adherent immune cells constitute a previously underappreciated conditioning layer that promotes biofilm formation on otherwise antifouling biomaterials. Pre-exposure of clinically used substrates to macrophages, monocytes, neutrophils or human peripheral blood cells markedly increased *Staphylococcus aureus* and *Escherichia coli* adhesion and aggregation. These studies also reveal that immune cells promote biofilms even after cell death with cellular debris acting as a conditioning agent. We demonstrate that reactive-oxygen-species amplification by incorporating bismuth telluride into a silicone composite converts adherent immune cells from passive conditioning agents into active bactericidal effectors. We note that this antimicrobial composite confers durable antibacterial protection across early, delayed and late infection time points in a murine implantation model. Together, these findings introduce a class of immune-coupled antibacterial materials as an alternative to the current antifouling paradigm.

## Introduction

Biofilm formation on indwelling medical devices is among the most intractable problems in healthcare. Bacterial colonization of catheters, prosthetic joints, vascular grafts and wound dressings drives a substantial fraction of healthcare-associated infections, frequently mandates device explantation, and remains a leading contributor to morbidity, mortality and cost worldwide^1^. Once established on a device surface, bacterial biofilms tolerate antimicrobials at concentrations several orders of magnitude above the planktonic minimum inhibitory concentration and resist clearance by host immune effectors^1^.

The most common strategy to overcome device-associated biofilms has been to modify the chemistry and topology of material surfaces. Examples of these efforts include the development and use of silicone and polytetrafluoroethylene as substrates for implants^2–4^, anti-adhesive chemistries based on poly (ethylene glycol), zwitterions and polysaccharide brushes^5,6^, antimicrobial strategies including silver ions and controlled release of small molecules from surfaces^5,7^, and nano/micro-topographical patterning to prevent microbe attachment^8–11^. These strategies reduce bacterial attachment substantially in controlled laboratory settings^8^. Despite these advances, the clinical incidence of device-associated biofilms remains a major challenge^1,7,12,13^. The gap between *in vitro* antifouling performance and clinical efficacy is now recognized, but its mechanistic basis is not understood.

A framework for understanding this gap was first proposed by Gristina as the “race for the surface^14^”, which states that the eventual fate of a biomaterial depends on whether host components or bacteria first colonize and condition the implant surface. Much of the research that followed has focused on specific proteins in this conditioning layer that serve as ligands to promote bacterial adhesion^15–18^. In contrast, the contribution of cellular conditioning to bacterial adhesion and biofilm formation remains poorly characterized^19,20^.

Most *in vitro* assays challenge pristine (or in a few cases protein-coated) materials directly with bacteria, but few interrogate surfaces that have previously been colonized by immune cells. This omission has direct clinical consequence. While bacteria may be introduced during implantation surgery, the dominant routes of late (days or weeks post implantation) device-associated infection are hub and lumen contamination, microbial tracking along percutaneous catheters, and hematogenous routes^21,22^. But, by this late time point, the device surface is no longer pristine, and it carries an established population of adherent immune cells and their debris^23,24^. The effect of this cellular conditioning layer on bacterial adhesion and its effects on engineered antifouling coatings has not been systematically evaluated.

In this study, we evaluate a hypothesis that adherent immune cells and immune-derived components generate a permissive substrate for bacterial adhesion and biofilm formation. To test this hypothesis, we exposed clinically used biomaterials to macrophages, monocytic and myeloid cell lines, and primary human immune cells prior to bacterial challenge. We compared the resulting bacterial adhesion, aggregation and biofilm formation against simultaneous (bacteria and immune cells added together) and bacteria-only controls. We also evaluated the durability of hyaluronic acid–poly(arginine) anti-bacterial coatings under each exposure sequence. We then asked whether bismuth telluride (Bi_2_Te_3_), a thermocatalytic material reported to modulate immune-cell behaviour and amplify localized reactive oxygen species generation^25^, could mitigate immune-cell-mediated biofilm potentiation when incorporated into a silicone composite. Finally, we validated our findings *in vivo* using a murine subcutaneous implantation model in which infection was deliberately staggered relative to implantation, allowing immune cell density at the implant surface to vary across infection time points. Our results identify cellular conditioning by adherent immune cells as a previously underappreciated determinant of biofilm formation on antifouling biomaterials, demonstrate that this effect is conserved across multiple cell types, bacterial species and material chemistries, and establish that converting adherent immune cells from passive conditioning agents into active bactericidal effectors by incorporating Bi_2_Te_3_ in the surface coating, is a temporally durable antibacterial strategy.

## Results

### Macrophage-biomaterial interactions promote *S. aureus* aggregation

To test the hypothesis that exposure to immune cells such as macrophages promote bacterial attachment and biofilm formation, we first used an *in vitro* setup where bacteria are added onto biomaterials without macrophages, simultaneously with macrophages, or following macrophage-biomaterial co-incubation for 12 hours, labelled as sequential cultures (**Figure 1a**). We chose poly(vinyl acetate) (PVAc) scaffolds that are used clinically for wound dressings^26,27^. For the initial studies, we chose *Staphylococcus aureus* (*S. aureus*) as it is commonly associated with implant-associated biofilm formation^28^, and specifically a clinically relevant *S. aureus* strain, ST88, which is known to aggregate and form biofilms^29,30^. In these experiments we observed that *S. aureus* monocultures exposed to the biomaterial alone showed some bacterial adhesion on material surface, but they did not exhibit aggregation (**Figure 1b and 1c**). In contrast, when the *S. aureus* was added simultaneously or sequentially with macrophages pronounced bacterial aggregation was observed (**Figure 1b and 1c**).

**Figure 1.**
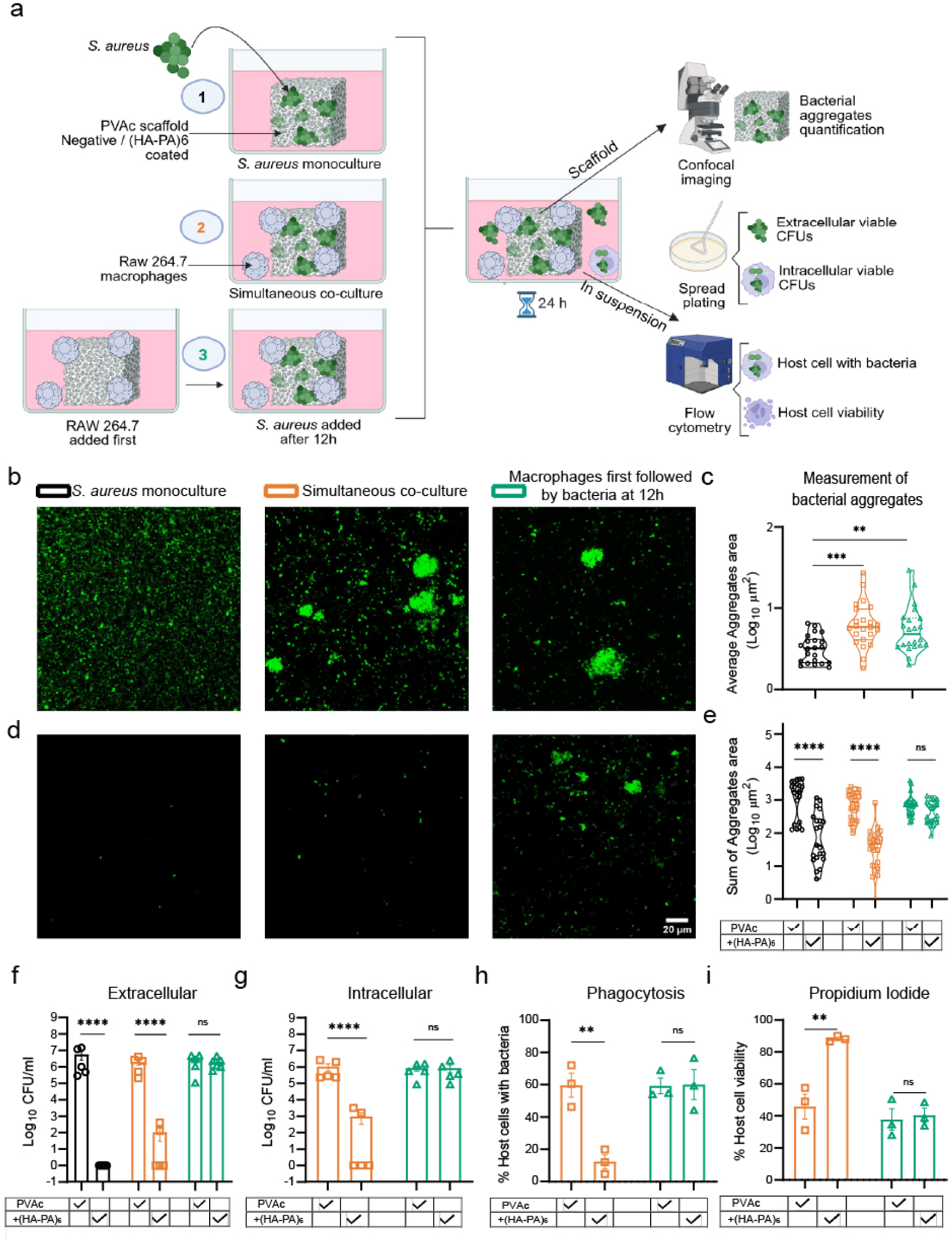
Macrophage-biomaterial interaction promotes bacterial aggregation on surfaces. (a) Schematic illustration of the experimental workflow showcasing the three different conditions for testing the hypothesis. Condition 1 (data presented as black circles) – no macrophages (control); condition 2 (data presented as orange squares) – simultaneous addition of macrophages and *S. aureus*; condition 3 (data presented as green triangles) – sequential addition of macrophages followed by *S. aureus*. (b) Representative confocal microscopy images of GFP-expressing *S. aureus* on biomaterial (PVAc) surfaces under different culture conditions. (c) Quantification of images from ‘b’, where the average area of bacterial aggregates is presented at different culture conditions; data is based on four independent experiments with minimum 5 fields of view (FOVs) analyzed per biomaterial. (d) Representative confocal microscopy images of GFP-expressing *S. aureus* on HA-PA coated PVAc surfaces under different culture conditions. (e) Quantification of images from ‘b’ and ‘d’, where total bacterial aggregate area (sum of all GFP signals) is presented at various culture conditions; data is based on four independent experiments with minimum 5 fields of view (FOVs) analyzed per biomaterial. (f) Assessment of planktonic extracellular surviving *S. aureus* in the supernatant expressed as colony forming units (CFU). (g) Evaluation of macrophage-internalized surviving *S. aureus* expressed as CFU. (h) Flow cytometric quantification of the percentage of live macrophages with GFP-expressing *S. aureus*. (i) Quantification of the percentage of live macrophages following infection with *S. aureus* under different treatment conditions. All CFU and flow cytometry results were presented as mean ± SD (n=5 for CFU; n=3 for flow cytometry). Statistical analysis was performed using one way ANOVA followed by Dunnett’s multiple comparison test, and two-way Anova with Sidak’s multiple comparison test. Statistical significance is indicated as \**p<*0.05, \*\**p<*0.01, \*\*\**p<*0.001, \*\*\*\**p<*0.0001, and ns=non-significant.

Surface modification is a commonly employed strategy to prevent bacterial adhesion and biofilm formation^31^. Hence, we next investigated whether surface functionalization using hyaluronic acid/poly(arginine) (HA–PA)^32^, enhances bacterial clearance. We first confirmed the antibacterial activity of poly(arginine) (PA) and HA-PA (where HA acts as the carrier for PA) have antibacterial efficacy against both *S. aureus* and *E. coli* under free-culture conditions (**Figure S1**). Subsequently, PVAc scaffolds were dip-coated with six alternating layers of HA and PA, and the HA–PA-coated PVAc scaffolds were confirmed to be non-cytotoxic to macrophages (**Figure S2**). We then evaluated these surface modified biomaterials under three defined culture conditions and observed that the HA–PA coating effectively reduced bacterial burden when macrophages were absent and when macrophages and bacteria were introduced simultaneously. However, when the biomaterial was preincubated with macrophages prior to bacterial challenge (sequential), the antibacterial efficacy of the coating was markedly diminished (**Figure 1d and 1e**).

We also measured the number of planktonic bacteria in the supernatant media and those internalized within macrophages and found both to be lower in HA-PA coated scaffolds compared to the uncoated scaffolds when no macrophages were present or macrophages and bacteria were added simultaneously. However, this effect was lost when the biomaterial was preincubated with macrophages prior to bacterial exposure (**Figure 1f and 1g**). The higher bacterial burden also resulted in worse outcomes for the macrophages in culture, with larger proportion of the cells showing intracellular bacterial presence (**Figure 1h**) resulting in lower viability (**Figure 1i)** in sequential but not simultaneous incubation conditions.

Collectively, these findings suggest that macrophages not only promote *S. aureus* aggregation on biomaterial surfaces but also compromise the antibacterial performance of surface-bound HA–PA coatings possibly by physically obstructing the interaction between bacteria and the PA coating.

### Silicone coating inhibits *S. aureus* aggregation, but high macrophage levels limit this inhibitory activity

Next, we asked if silicone, a more commonly used material in medical implants and indwelling catheters, would help mitigate these effects of macrophage-biomaterial interaction induced bacterial adhesion and aggregation. We evaluated the antibacterial performance of silicone coatings alone and in combination with HA–PA under the sequential exposure conditions described in figure 1a. As would be expected based on previous reports, silicone coating resulted in a significant decrease in bacterial adhesion and aggregation, which was further lowered when HA-PA was incorporated on silicone-coated surfaces (**Figure 2a and 2b**). As another validation of these observations, bacterial aggregates were recovered from biomaterial surfaces via bath sonication, and viable bacteria were enumerated using a colony forming unit (CFU) analysis, which showed that silicone and silicone-HA-PA resulted in approximately half-log-fold lower number of viable bacteria (**Figure 2c**). In parallel, bacterial burden in the culture supernatant and within macrophages was seen to be significantly lower in the silicone-HA-PA but not the silicone coated surfaces (**Figure S3a-d**). These results suggested that silicone, which is known to prevent bacterial adhesion due to its hydrophobic nature, does lower bacterial attachment and aggregation, and this may be further improved with a HA-PA coating on the silicone surface.

**Figure 2.**
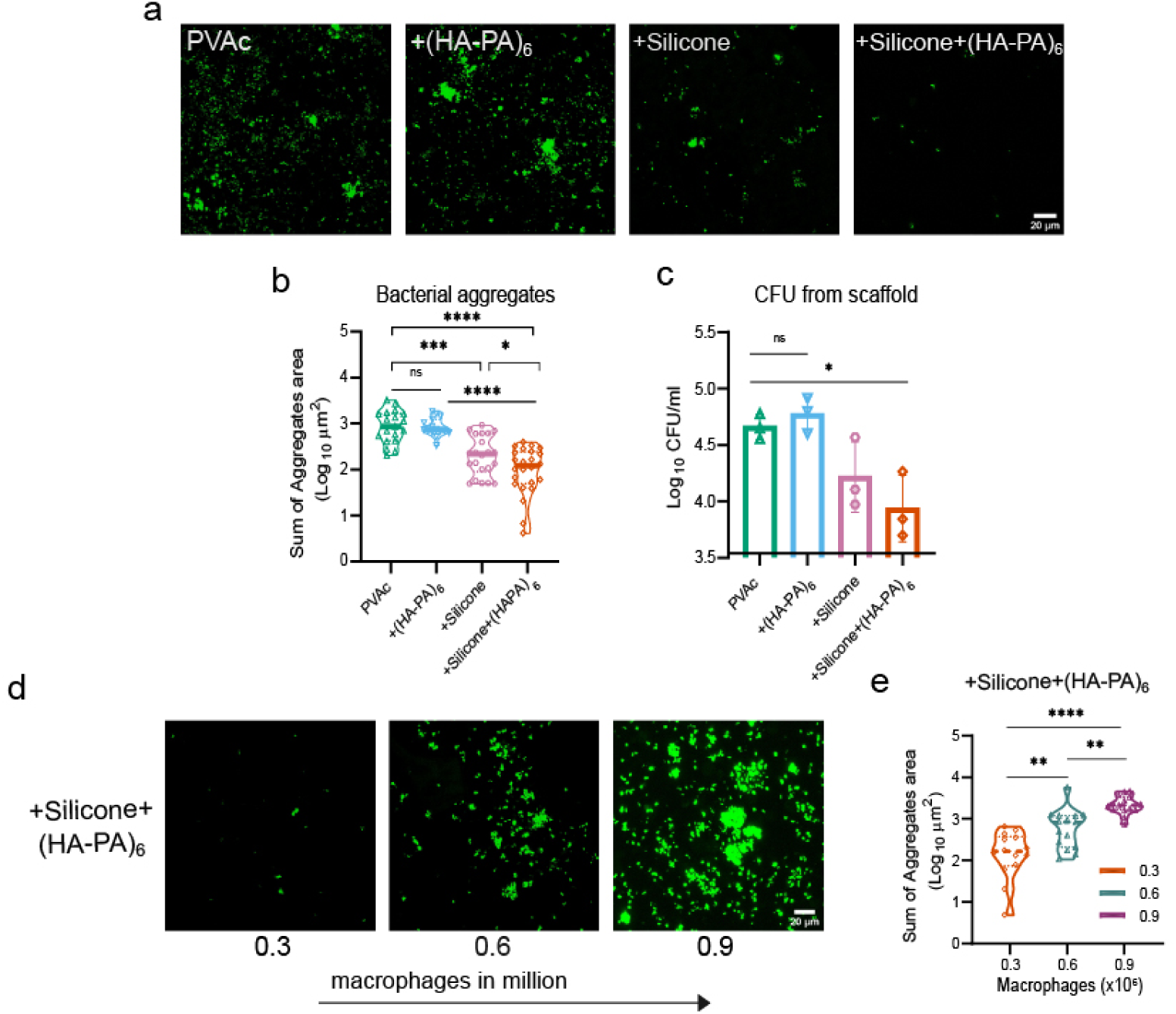
Effect of silicone and silicone-HA-PA coating on *S. aureus* aggregation on the surface of the material. (a) Representative confocal microscopy images of GFP-expressing *S. aureus* on biomaterial surfaces in the presence of different coatings (PVAc is the base material). (b) Quantification of images presented in ‘a’ as total bacterial aggregation on the biomaterial surfaces with different coatings. (c) Quantification of surviving *S. aureus* recovered from the biomaterial, reported as CFU. (d) Representative confocal microscopy images of GFP-expressing *S. aureus* Silicone+HA-PA coated surfaces in the sequential experimental condition (figure 1a) where macrophage numbers were varied as 0.3, 0.6 or 0.9 million cells per scaffold. (e) Quantification of images presented in ‘d’, where sum of bacterial aggregate area is presented. Bacterial aggregate quantification was performed using data from four independent experiments with minimum 4 FOVs analyzed per biomaterial. Statistical analysis was performed using one way ANOVA followed by Tukey’s multiple comparison test. Statistical significance is indicated as *p<0.05, **p<0.01, ***p<0.001, ***p<0.0001, and ns=nonsignificant.

To challenge the silicone-HA-PA coated system further, we examined if increasing the macrophage number would affect the ability of the silicone surfaces to reduce bacterial attachment and aggregation. Increasing macrophage numbers from 0.3 million to 0.9 million (per scaffold) resulted in higher bacterial adhesion and aggregation (**Figure 2d and 2e**). These data suggest that: (i) the silicone-HA-PA coated system is liable to macrophage-biomaterial interaction-based increase in bacterial adhesion and aggregation, and that (ii) a larger number of macrophages are required to facilitate this effect.

We confirmed that the finding of increasing macrophage number promoting bacterial adhesion and aggregation was also seen on PVAc surfaces (**Figure S4a and b**), but that a very high number of macrophages (3 million) could prevent the effect possibly through direct macrophage-mediated bacterial clearance under these conditions (**Figure S4a and c**). To probe bacterial phenotypic changes associated with macrophage-induced aggregation, bacteria recovered from biomaterial surfaces were plated on sheep blood agar. We observed a pronounced reduction in zones of hemolysis (**Figure S4d**), indicative of decreased δ-hemolysin activity and agr expression, consistent with repression of agr-regulated biofilm-disrupting factors^33^. These observations suggest that enhanced interactions between macrophages and bacteria in the biomaterial microenvironment induce transcriptional reprogramming in *S. aureus*, shifting bacterial behavior toward a biofilm-dominant, aggregation-prone state. This finding underscores the critical importance of regulating immune cell–bacteria interactions at biomaterial interfaces to prevent biofilm establishment.

### Immune cell-biomaterial interaction mediated bacterial aggregation and biofilm formation holds true across bacterial species, biomaterials and cell types

A question of interest was whether these observations would hold true for other bacterial species, biomaterials and cell types. To address this question, we evaluated the dominant human bacterial pathogen *Escherichia coli* (*E. coli*) as the bacterial species, silicone coated on polystyrene tubes resembling indwelling catheters as the biomaterial with the polystyrene tubes posing as additional material surfaces, and THP-1 monocytes, HL-60 myeloid cells, and human blood cells as the cell types. We assessed the bacterial aggregation and biofilm formation using a crystal violet assay. Even with these changes, we observed a clear and significant increase in biofilm formation upon the sequential addition of immune cells followed by bacteria (**Figure 3a-c**), suggesting that immune cell adhesion on a material surface drives bacterial aggregation and biofilm formation.

**Figure 3.**
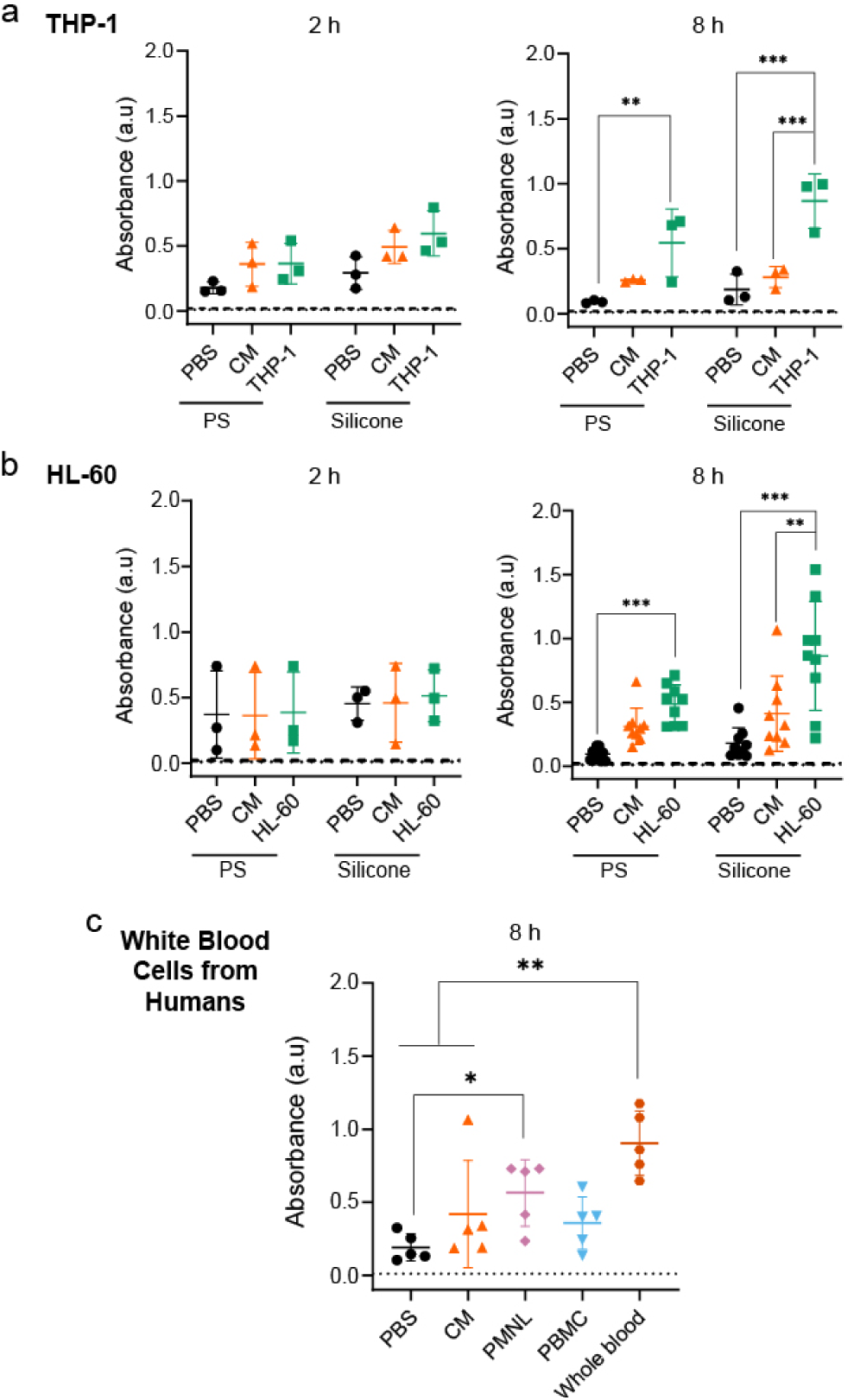
Immune cell-biomaterial interaction involving THP-1, HL-60 cell lines and peripheral venous white blood cells from humans also demonstrate increased bacterial biofilm formation. (a) Biofilms were quantified using a crystal violet assay on tubes of polystyrene (PS) or polystyrene coated with silicone (Silicone) exposed to either phosphate buffered saline (PBS), complete media (CM) or THP-1 cells in complete media for 2 and 8 hours, followed by *E. coli* incubation for 48 hours. (b) and (c) similar experimental setup as ‘a’, with the cells changes to HL-60 (b) or various cells present in human blood (c), where PMNLs refers to polymorphonuclear leukocytes (PMNLs) and PBMCs refers to peripheral blood mononuclear cells (PBMCs). In (c) polystyrene coated with silicone was used. One-way ANOVA followed by Tukey’s multiple comparisons test was used for statistical comparison. Dotted line/dashed line in the graphs indicate crystal violet absorbance of bare polystyrene/silicone tube (background absorbance). *p<0.05, **p<0.01, ***p<0.001.

### Bacterial aggregates spatially colocalize with dead macrophages, and Bismuth Telluride mitigates this effect

Next, we examined where the bacterial aggregates were forming on the biomaterial surface. Using the PVAc as the base material along with its surface modifications, and through confocal imaging of GFP-tagged *S. aureus* along with propidium iodide (PI) staining to identify dying or dead macrophages, we assessed the co-localization of the macrophages and bacteria. Strikingly, we observed that a substantial proportion of the *S. aureus* aggregates were spatially colocalized with dead macrophages, an effect that was markedly amplified under high macrophage (0.9 million cells) density conditions (**Figure 4a**). To validate that host cell death contributes to bacterial aggregation and biofilm formation, macrophages were heat-inactivated by incubation at 60□°C for 30 minutes. The resulting lysed cellular components were then incubated with silicone-coated PVAc surfaces. Notably, *S. aureus* aggregation was observed in the presence of heat-killed macrophages. This observation was corroborated using HL-60 cells and *E. coli* bacteria, where components from lysed (H_2_O_2_ based lysis) cells on silicone-coated polystyrene tubes led to a similar increase in biofilm formation **(Figure S5)**.

**Figure 4.**
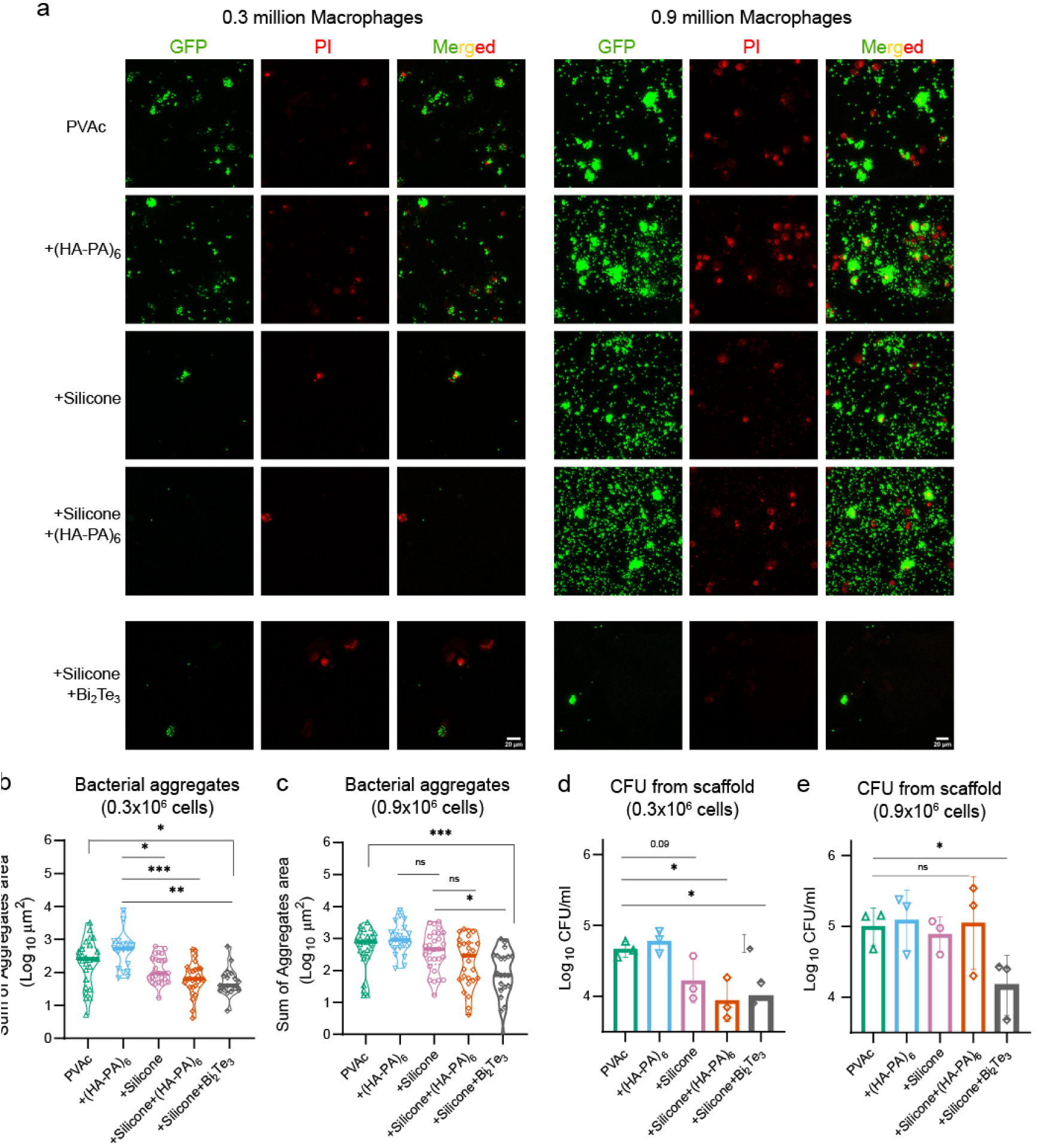
Spatial association of dead macrophages with *S. aureus* bacterial aggregates and bismuth telluride reduces bacterial aggregate on material surfaces. (a) Representative confocal images of biomaterial surfaces stained with propidium iodide to label membrane-compromised macrophage nuclei (red) and GFP-expressing *S. aureus* (green) under different coating conditions at macrophage densities of 0.3 and 0.9 million. (b) Representative confocal images with bismuth telluride (Bi_2_Te_3_) coated silicone-PVAc surfaces at macrophage densities of 0.3 and 0.9 million. (c) and (d) Quantitative analysis of total bacterial aggregates area at macrophage densities of 0.3 million (c) and 0.9 million (d). (e) and (f) Quantification of surviving S. aureus CFU recovered from the biomaterial at macrophage densities of 0.3 million (e) and 0.9 million (f), reported as CFU. Data is presented as mean ± SD (n=3 independent experiments); Bacterial aggregate quantification was performed using data from three independent experiments with minimum 4 FOVs analyzed per biomaterial. Statistical analysis was performed using one way ANOVA followed by Tukey’s multiple comparison test. Statistical significance is indicated as *p<0.05, **p<0.01, ***p<0.001, and ns=non-significant.

To mitigate these bacterial aggregation and biofilm formation events driven by macrophage-biomaterial interactions, we explored the possibility of using bismuth telluride as a coating as it demonstrates thermoelectric and thermo-catalytic effects that can modulate macrophage adhesion and possibly enhance ROS production of the whole system^25,34,35^.

First, bismuth telluride (Bi_2_Te_3_) hexagonal nanostructures were prepared and TEM characterization showed defined hexagonal nanosheets with lateral dimensions in the sub-micrometer range, indicating controlled crystal growth and high structural uniformity (**Figure S6a**). The corresponding selected area electron diffraction (SAED) pattern displayed sharp and periodically arranged diffraction spots, confirming the highly crystalline nature of material, and the crystal structure revealed layered atomic arrangement of Bi_2_Te_3_ nanoflakes (**Figure S6b-c**). This highly ordered crystallinity is important for efficient carrier transport and minimizing charge recombination during catalytic operation. The thermal catalytic performance of Bi_2_Te_3_ was evaluated under different temperature differences (ΔT = 7, 10, and 13 °C) and showed gradual increase in H_2_O_2_ generation (**Figure S6d**). The proposed thermally driven catalytic mechanism is illustrated in **figure S6e**. Under thermal stimulation, a temperature gradient could be established across the nanostructure, producing spatial charge separation between hot and cold regions wherein electrons accumulate on one side of the material surface, while positive charges remain on the opposite side. This asymmetric charge distribution creates an internal electric field that promotes interfacial redox reactions. The accumulated electrons react with dissolved oxygen to generate superoxide intermediates (•O_2_⁻), which are subsequently converted into H_2_O_2_. Further, Kelvin probe force microscopy (KPFM) measurements support temperature-dependent charge generation of the material surface with surface potential progressively increasing from 195 mV at ΔT = 0 °C to 241 mV at ΔT = 13 °C (**Figure S6f-i**). Together, these results demonstrate the ability of Bi_2_Te_3_ to convert low-grade thermal energy into catalytic activity, and hence their potential for use as part of biomaterials to reduce bacterial adhesion.

The Bi_2_Te_3_ nanoflakes were incorporated into a silicone matrix and applied as a composite coating on the PVAc surface. Initial cytocompatibility assessments confirmed that the bismuth telluride–silicone coating was non-cytotoxic toward macrophages (**Figure S7a**). Notably, under both low and high macrophage density conditions (0.3 and 0.9 million cells), the composite coating of Bi_2_Te_3_ and silicone resulted in a pronounced suppression of bacterial aggregation, as evidenced by both confocal microscopy (**Figure 4c and 4d**) and viable CFU enumeration (**Figure 4e and 4f**).

To evaluate the broader immunological relevance of this effect, we extended our analysis to primary human immune cells. While human PBMCs and neutrophils similarly promoted bacterial aggregation on unmodified biomaterials, the Bi_2_Te_3_–silicone coating consistently exhibited superior antibacterial efficacy across both low and high immune cell densities (**Figure S7b-i**). Importantly, this enhanced performance was not limited to *S. aureus*. Indeed, against *E. coli*, the Bi_2_Te_3_–silicone coating significantly reduced bacterial burden under both cell-free and immune cell–rich conditions, whereas previously tested coatings failed to achieve comparable efficacy (**Figure S8**).

### Bismuth telluride enhances macrophage-mediated bactericidal activity *against S. aureus* through the induction of reactive oxygen species

In planktonic, cell-free bacterial cultures, Bi_2_Te_3_ exhibited no antibacterial activity against *S. aureus* and minimal antibacterial activity against *E. coli* (**Figure S9**). This observation was initially counterintuitive, given the pronounced suppression of bacterial aggregation at the Bi_2_Te_3_-silicone biomaterial interface observed in the presence of immune cells. These findings prompted us to hypothesize that Bi_2_Te_3_ may not function as a direct antibacterial agent, but instead it potentiates immune cell-mediated bactericidal mechanisms through increased production of reactive oxygen species (ROS). To test this hypothesis, bacterial burden was quantified by CFU enumeration under cell-free conditions and in coculture with immune cells, with and without Bi_2_Te_3_. Bi_2_Te_3_ treatment did not result in a statistically significant reduction in planktonic extracellular bacteria in the absence of immune cells, but in the presence of immune cells, Bi_2_Te_3_ induced a marked reduction in both extracellular planktonic bacteria and intracellular bacterial populations (**Figure 5a-c**), indicating an immune-dependent antibacterial effect. We next assessed ROS generation by quantifying superoxide levels in culture supernatants in the absence of bacteria. Notably, superoxide concentration was increased only in the presence of immune cells (**Figure 5d**). These data indicate that Bi_2_Te_3_ leverages the increased metabolic and oxidative potential of larger immune cell populations to amplify ROS-mediated bacterial killing. To further validate these observations, biomaterial coated with Silicone+Bi_2_Te_3_ were exposed to heat-killed macrophages in place of live cells. Under these conditions, the previously observed antibacterial activity of Bi_2_Te_3_ was completely abrogated. This loss of efficacy in the presence of non-viable, lysed cellular components strongly suggests that the antibacterial effect is contingent upon interactions with live host cells (**Figure S10**).

**Figure 5.**
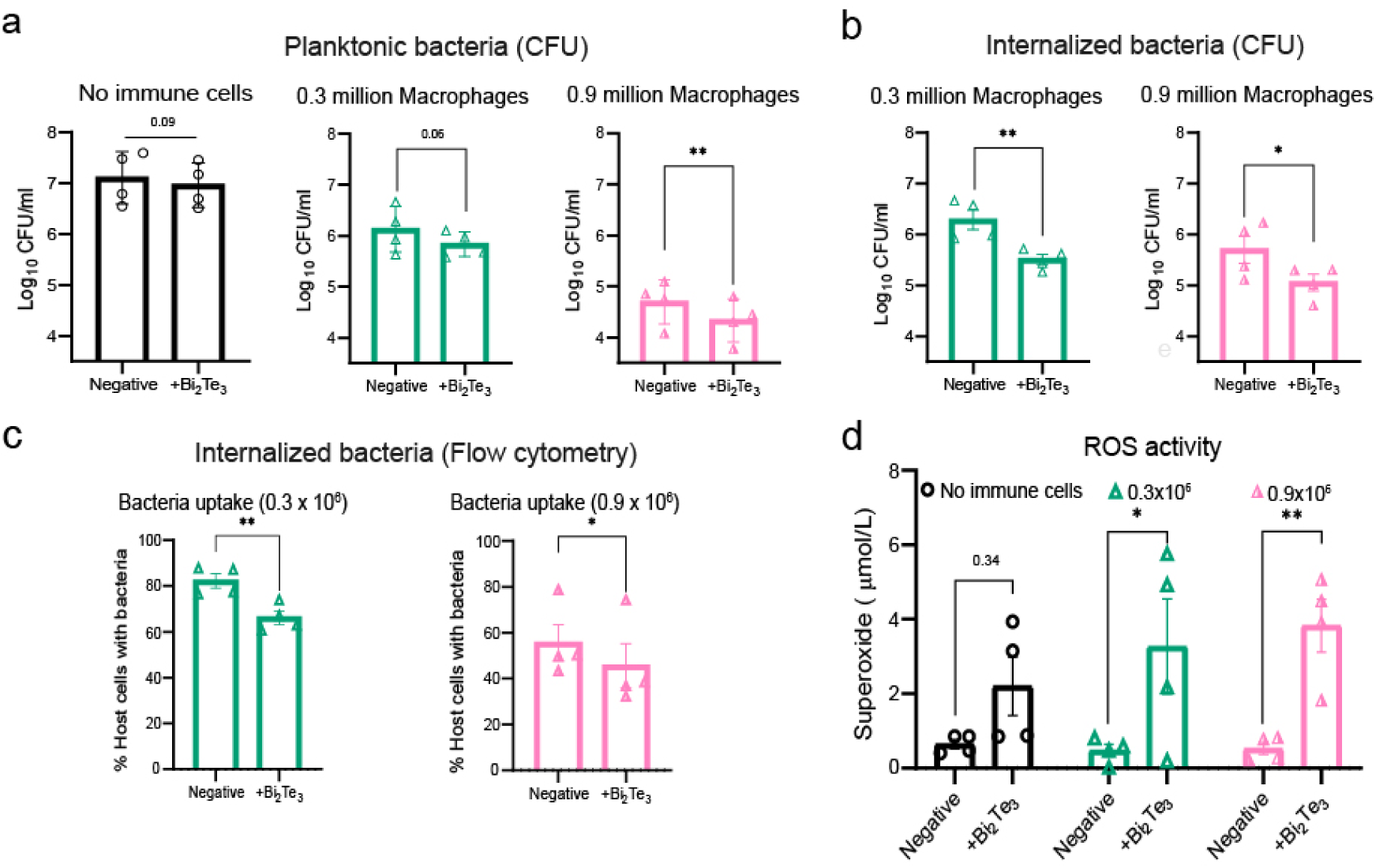
Bismuth telluride enhances macrophage-mediated bactericidal activity *against S. aureus* through the induction of reactive oxygen species. (a) Quantification of extracellular planktonic surviving *S. aureus* in the supernatant of cultures in the absence and presence of macrophages at densities of 0.3 million and 0.9 million expressed as CFU; negative indicates absence of bismuth telluride (Bi_2_Te_3_) and + Bi_2_Te_3_ indicates a dose of 0.3 mg bismuth telluride added to these cultures. (b) Evaluation of internalized surviving *S. aureus* recovered from macrophages in coculture at macrophage densities of 0.3 million cells and 0.9 million cells, in the absence and presence of bismuth telluride; number of bacteria is expressed as CFU. (c) Flow cytometric quantification of the percentage of live macrophages associated with GFP-expressing *S. aureus* (d) Quantitative analysis of superoxide production by macrophages in the absence (negative) or presence of bismuth telluride (Bi_2_Te_3_). No bacteria were presented in these experiments. Data are presented as mean ± SD (n=4 independent experiments); Statistical analysis was performed using paired t-test and two-way ANOVA followed by Sidak’s multiple comparison test. Statistical significance is indicated as *p<0.05, **p<0.01, ***p<0.001, ****p<0.0001, and ns=non-significant.

### *In-vivo* implant-biofilm model demonstrates that Bi_2_Te_3_ confers sustained anti-biofilm protection

To validate our *in vitro* findings under physiologically relevant conditions, we next evaluated the antibacterial performance of the various coated biomaterials *in vivo* using a murine biomaterial implantation model. Biomaterials bearing different surface modifications were implanted subcutaneously (marked as day 0), while bacterial challenge with *S. aureus* was administered either immediately (day 0), after 5 days, or after 12 days, representing early, delayed, and late infection scenarios, respectively (**Figure 6a**). These staggered infection time points were designed to progressively increase immune cell presence on the biomaterial surface, and to recapitulate clinically relevant infection routes, namely intraoperative contamination, delayed post-surgical infection due to impaired wound healing, and late hematogenous or wound-associated infection^22^. Although the timing of infection varied, the infection duration was maintained at 24 hours across all conditions. Biomaterials were explanted the day following infection and assessed for biofilm formation using scanning electron microscopy and confocal imaging.

**Figure 6.**
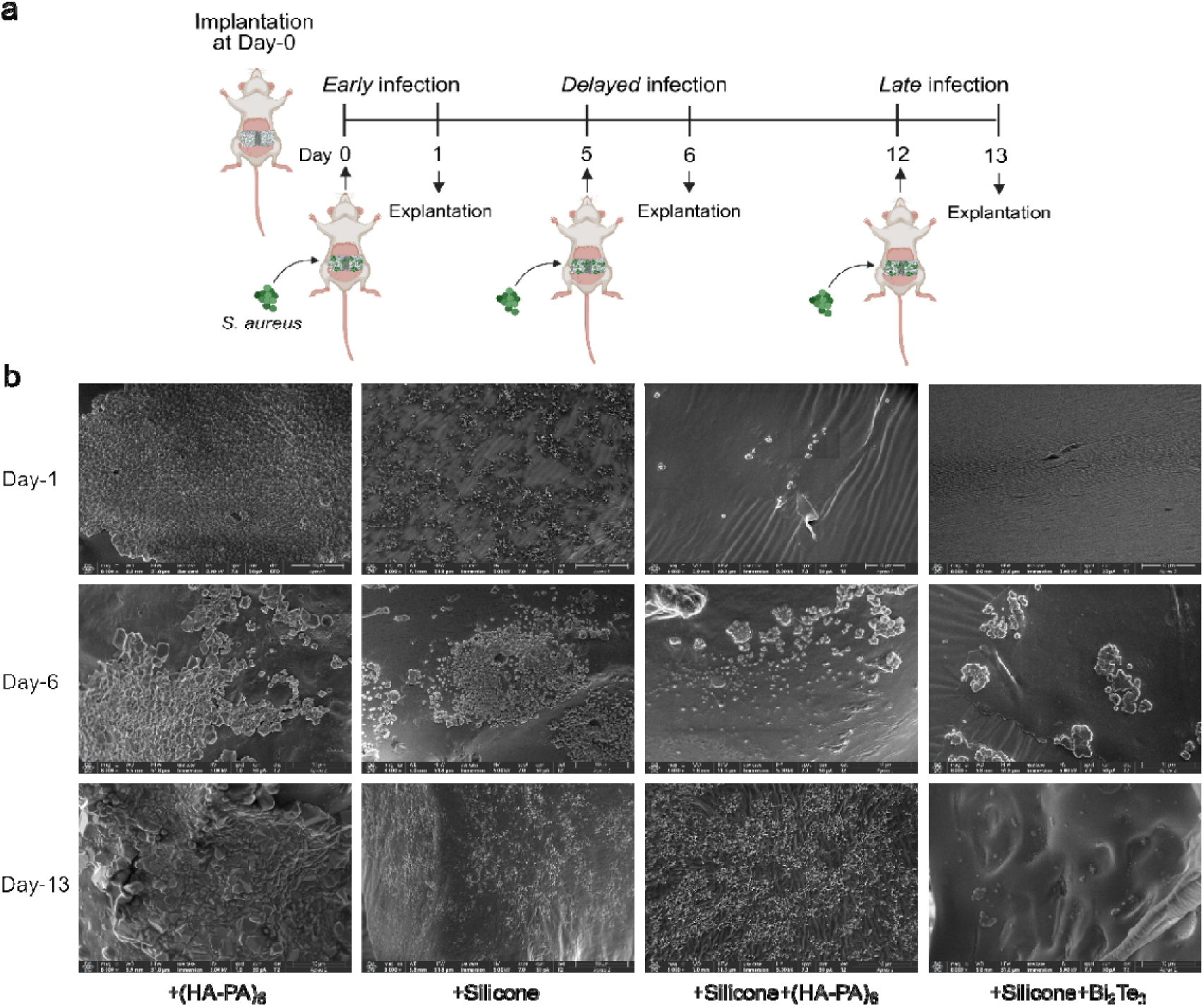
*In vivo* validation reveals that Silicone and HA-PA coatings effectively limit biofilm only during early infection, whereas bismuth telluride retains anti-biofilm efficacy against *S. aureus* even under conditions of late infection. (a) Schematic showing the design of the *in vivo* experiments, where bacteria were injected during the biomaterial implantation surgery (early xinfection), or at 5-days post-surgery (delayed infection), or at 12-days post-surgery (late infection); biomaterials were retrieved 1 day post bacterial infection in all conditions. (b) Representative scanning electron micrographs illustrating bacterial adhesion and biofilm formation on implanted biomaterials at early, delayed and late infection time points

At the early (day 0) infection time point, uncoated and HA–PA-coated biomaterial exhibited substantial biofilm formation, whereas silicone-based coatings, including silicone alone, silicone–HA–PA, and silicone–bismuth telluride (Si–Bi_2_Te_3_), effectively suppressed bacterial colonization (**Figures 6b, 7a and 7b**). By the delayed infection stage of 5 days post implantation, silicone alone began to show measurable biofilm accumulation, while both silicone–HA–PA and silicone– Bi_2_Te_3_ coatings continued to limit bacterial aggregation.

**Figure 7.**
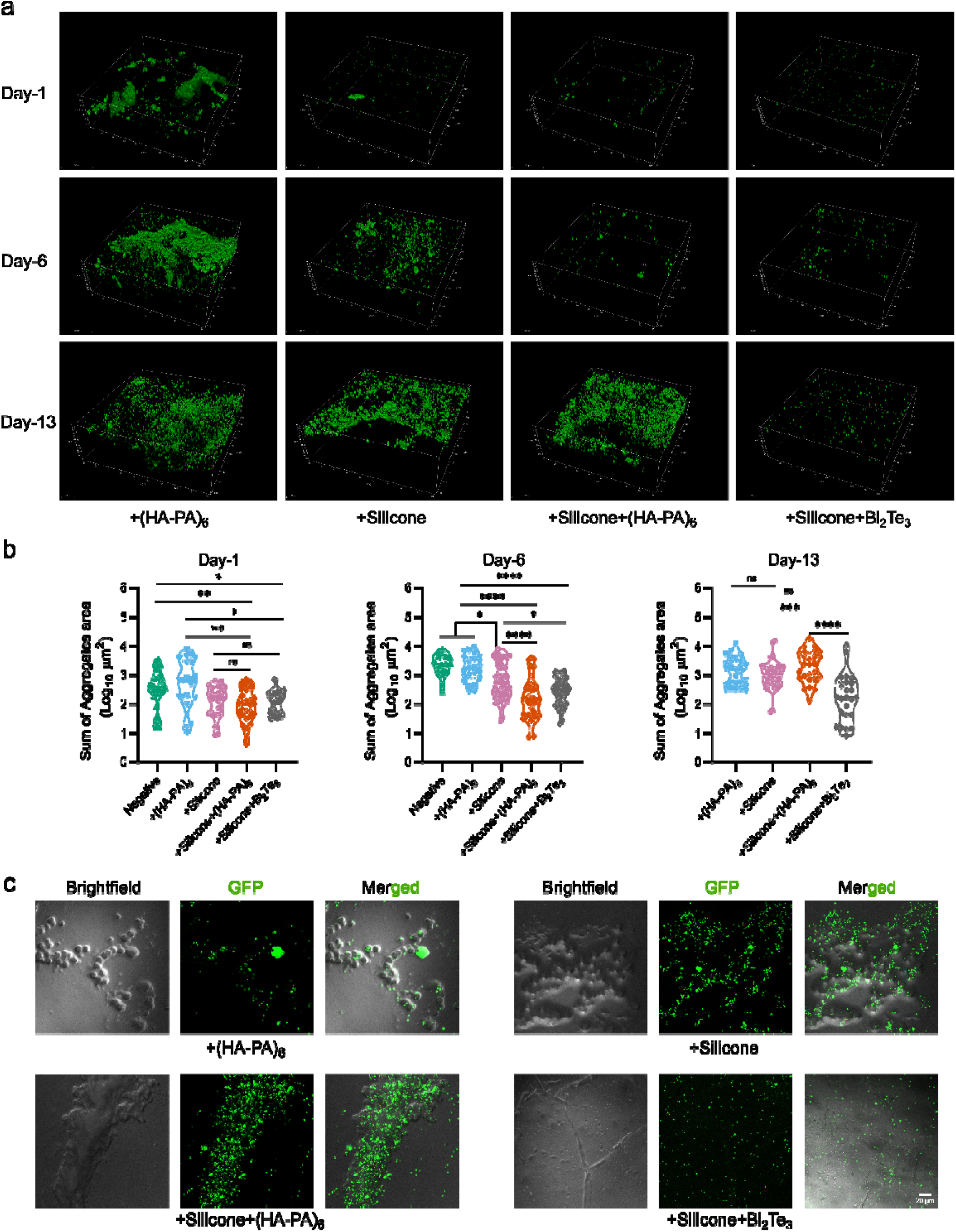
Confocal imaging of explanted biomaterials reveals that bismuth telluride maintains reduced immune cell adhesion and robust anti-biofilm efficacy. (a) Representative confocal images (3D representation) of explanted biomaterials illustrating GFP-expressing *S. aureus* mediated biofilm formation on surfaces at early, delayed, and late infection time points. (b) Quantitative analysis of images presented in ‘a’, showing total bacterial aggregate area (calculated as sum of GFP fluorescence intensity) across different coating conditions and explanted time points. (c) Representative images (∼12 of 24 fields of view (FOVs) of each biomaterial surface except for silicone-Bi2Te3 which is based on 22 of 24 FoVs) showing immune cells (brightfield) and bacterial aggregates on explanted biomaterials at day-13. Bacterial aggregate quantification was performed with a minimum of 6 FOVs analyzed per biomaterial (at least 4 repeats for each biomaterial). Statistical analysis was performed using one way ANOVA followed by Tukey’s multiple comparison test. Statistical significance is denoted as \**p<*0.05, \*\**p<*0.01, \*\*\**p<*0.001, \*\*\*\**p<*0.0001, and ns=non-significant.

Significantly, during late-stage infection (12 days post implantation), while both silicone and silicone–HA–PA coatings failed to prevent biofilm formation, the silicone–Bi_2_Te_3_ coating retained robust anti-biofilm efficacy, exhibiting a marked reduction in bacterial burden relative to all other surface modifications (**Figure 7a and 7b**). Importantly, spatial analysis revealed frequent colocalization of immune cell clusters with bacterial aggregates at the biomaterial interface in the late-stage (12-days post implantation) infection setting for all coating conditions, except for silicone–Bi_2_Te_3_ (**Figure 7c**). The colocalization was consistently observed in at least 12 of 24 fields of view per biomaterial for the +HA-PA, +silicone, and +silicone-HA-PA coated surfaces. In striking contrast, immune cell–bacteria colocalization events were rarely detected on silicone–Bi_2_Te_3_ coated biomaterials, with only two occurrences observed across 24 fields of view from four biomaterials (**Figure 7c**). These *in vivo* observations further corroborate our mechanistic findings that immune cells play an active role in promoting biofilm formation under macrophage-rich conditions. The anti-bacterial performance of silicone–Bi_2_Te_3_ coatings suggests that incorporation of bismuth telluride not only prevents bacterial aggregation but also mitigates immune-cell-mediated biofilm stabilization, likely by restoring or enhancing host bactericidal activity at the biomaterial interface. Collectively, these results establish silicone–Bi_2_Te_3_ as a uniquely durable antibacterial strategy capable of maintaining efficacy across the full temporal spectrum of infection.

## Discussion

In this study, we tested if adhesion of immune cells on biomaterials promoted bacterial colonization of the surface. Across five cell types (RAW 264.7 macrophages, THP-1 monocytes, HL-60 myeloid cells, and primary human PBMCs and neutrophils), two bacterial species (*S. aureus* and *E. coli*), three base materials (PVAc, silicone, polystyrene), and both *in vitro* and murine *in vivo* settings, we observed a consistent pattern that pre-conditioning of biomaterial surfaces with immune cells substantially increased bacterial adhesion, aggregation and biofilm formation. The effect was sufficient to abrogate the antibacterial activity of an HA–PA coating that performed well against simultaneous bacterial challenge, and partially overcome the antifouling behaviour of silicone. Incorporation of bismuth telluride (Bi_2_Te_3_) into a silicone matrix prevented the immune-cell mediated potentiation of bacterial biofilm formation, both *in vitro* across multiple immune cell types and *in vivo* across early, delayed and late infection time points, while exhibiting negligible direct antibacterial activity in cell-free conditions. These findings demonstrate that immune cell adhesion is a critical determinant of clinical antifouling failure, and identify immune-cell-coupled bactericidal materials as a distinct design strategy for indwelling medical devices.

Our observations extend Gristina’s race for the surface^14^ framework in a direction that has received comparatively little attention. The classical formulation envisions host cells and microbes as competitors for an initially pristine surface, with host victory promoting tissue integration and microbial victory yielding infection. Our data suggest that this dichotomy is inaccurate. That is, when host immune cells reach the surface first, they do not simply outcompete bacteria, rather, they could condition the interface in a manner that promotes subsequent bacterial colonization, particular when cellular viability is compromised. The spatial colocalization of *S. aureus* aggregates and propidium iodide–positive macrophages, both *in vitro* and at late infection time points *in vivo*, supports a model in which dying immune cells serve as a cellular conditioning layer, which may be thought of as an extension of the protein-corona conditioning concept^5,21^.

The mechanisms by which adherent immune cells potentiate bacterial colonization appear to involve the death of these cells, which may provide both nutrient substrates and extracellular DNA, the latter of which is a known structural component of the staphylococcal biofilm matrix^36^. The data involving lysed HL-60 cells promoting biofilm formation on silicone-coated tubes and the spatial colocalization of bacteria with dead cells on PVAc scaffolds suggests that this mechanism is likely to be true. However, our data do not exclude the possibility of alternate or additional supporting mechanisms, such as the secretion of soluble factors by cells that are alive and possibly stressed on the implant surface, or bacterial environmental sensing within the immune-cell-rich microenvironment.

The dose-response relationship between macrophage density and bacterial aggregation appeared to be non-monotonic. Low macrophage numbers (∼0.3 × 10^6^ / scaffold) produced limited conditioning and modest biofilm enhancement, intermediate densities (∼0.9 × 10^6^ / scaffold) produced the most pronounced potentiation, while very high densities (∼3 × 10^6^) suppressed biofilm formation presumably through direct phagocytic clearance outpacing surface conditioning. These data imply that biofilm enhancement is driven specifically by the regime in which cellular conditioning exceeds active bactericidal capacity, a regime that is possibly the steady-state cellular environment around chronic implants where a finite but non-overwhelming population of macrophages and foreign-body giant cells persist at the interface^23,24^. The temporal pattern of our *in vivo* findings, in which silicone and silicone–HA–PA coatings retained efficacy at early infection time points but failed by day 12 post-implantation, mirrored by progressive accumulation of immune cells at the implant surface, is consistent with this interpretation.

Bi_2_Te_3_ was selected as a candidate to overcome the limitations of other coatings based on prior reports that thermoelectric and thermo-catalytic chalcogenides can amplify reactive oxygen species generation in the presence of metabolically active cells^25,34,35^. Our data are consistent with such a mechanism, where Bi_2_Te_3_ exhibited negligible bactericidal activity in cell-free planktonic culture, but markedly reduced both extracellular and intracellular bacterial burden in the presence of immune cells, and elevated supernatant superoxide levels only when immune cells were present. We propose that Bi_2_Te_3_ functions not as a direct antimicrobial but as an immune-coupled bactericidal amplifier, converting adherent immune cells from passive conditioning agents into active killers at the device interface.

However, several aspects of the mechanism by which Bi_2_Te_3_ coatings might function remain to be defined. In this context, a mechanistic caveat warrants emphasis. The thermo-catalytic H_2_O_2_ generation we characterized for Bi_2_Te_3_ (**Figure S6**) requires an imposed temperature gradient, whereas the cell-culture and subcutaneous environments in which the coating proved effective are close to isothermal. Additionally, the benefit of Bi_2_Te_3_ was strictly dependent on the presence of viable immune cells and was accompanied by elevated supernatant superoxide only when cells were present, with no measurable bactericidal activity in cell-free planktonic culture (**Figure 5**). Taken together, these observations indicate that the operative mechanism at the biological interface is unlikely to be autonomous thermo-catalytic H_2_O_2_ production by the material. Rather, Bi_2_Te_3_ appears to act as an immune-coupled amplifier of cell-derived reactive oxygen species. We speculate that the relevant *in vivo* trigger could be a combination of subtle temperature gradient, redox potential, and cell-membrane contact, but the precise mechanism remains to be identified.

Another point of note is the difference between the conditioning (immune cell presence conditions material surfaces for bacterial biofilm formation) and the therapeutic (Bi_2_Te_3_ coating reduces bacterial biofilm formation) aspect of the study. Surface conditioning that potentiated biofilm formation was reproduced by non-viable cellular material, including heat-killed macrophages and lysed HL-60 cells, implying that dead cells and debris are sufficient to render a surface permissive to bacterial aggregation. In contrast, the bactericidal benefit of Bi_2_Te_3_ was abolished when live cells were replaced with heat-killed cells, indicating a requirement for metabolically active cells. These results are reconcilable if the chronic implant interface is viewed as a dynamic mixture of metabolically active (viable) immune cells and accumulating biomolecular and cellular debris. In this setting Bi_2_Te_3_ would act on the live fraction to shift the local balance from permissive conditioning toward active killing. This interpretation suggests that Bi_2_Te_3_ efficacy will depend on the ratio of viable to non-viable cells at the surface, a parameter that warrants direct testing in future work.

In conclusion, our data argue for a substantive revision of how antibacterial biomaterials are designed and evaluated. Standard antifouling assays performed against pristine surfaces are likely to overestimate clinical performance, and laboratory protocols that pre-condition surfaces with relevant immune cells before bacterial challenge would provide a more predictive readout. From a design standpoint, our results suggest that the most durable antibacterial strategies for indwelling devices may not be those that minimize host–material interaction altogether, which is an approach that has mostly failed to translate. Instead, a focus on materials that productively couple the inevitable host cellular response to bactericidal function could be the way forward.

## Materials and Methods

### Materials and Reagents

Medical grade polyvinyl acetate (PVAc) based Ivalon nasal packs were procured from Syntron Health Care (India). The murine macrophage cell line RAW 264.7 was procured from Sigma Aldrich, and the human promyelocytic neutrophil-like cell line HL-60 and human monocytic cell line THP-1 were obtained from the American Type Culture Collection (ATCC, USA). Hyaluronic acid (HA; M_w_: 25,000), poly-L-arginine hydrochloride (PA; M_w_: 5,000-15,000), chloramphenicol and ampicillin were purchased from Sigma-Aldrich, Merck, USA. Glass-bottom dishes (35 mm) were sourced from ibidi (Germany). Tryptone Soya broth (TSB) and Luria Bertani (LB) were obtained from HiMedia (India). Dulbecco’s Modified Eagle Medium (DMEM) was purchased from MP Biomedicals (USA), while Fetal Bovine Serum (FBS) and antibiotic–antimycotic solution (1%) were acquired from ThermoFisher Scientific (USA).

### Bacterial Strains and Growth Conditions

Community-acquired methicillin-resistant *Staphylococcus aureus* (CA-MRSA) ST88 strain harboring the pTH100 plasmid (Addgene, USA), which encodes the superfold green fluorescent protein (sGFP), was cultured in TSB supplemented with chloramphenicol (10□µg/mL) at 30□°C for 10□h with agitation at 180□rpm. Similarly, *Escherichia coli* DH10β carrying the pDiGc plasmid (Addgene, USA), enabling constitutive expression of enhanced green fluorescent protein (eGFP), was grown in LB containing ampicillin (100□µg/mL) under identical conditions (30□°C, 10□h, 180□rpm shaking). For experiments involving silicone coated polystyrene tubes, non-fluorescent *E. coli* strain MG1655 was cultured at similar conditions at 37□°C for 10□h with agitation at 180□rpm shaking.

### Immune Surveillance and In vitro Infection Assays

RAW 264.7 cells were maintained in DMEM supplemented with 10% FBS and 1% antibiotic–antimycotic solution. For immune surveillance and infection assays, PVAc strips (8 cm) were hydrated in phosphate-buffered saline (PBS; pH 7.4) to swell up, sterilized by autoclaving, and sectioned into scaffolds measuring approximately 4 × 4 × 2 mm^3^. These scaffolds were gently placed in 24-well plates containing RAW 264.7 cells in 400 µL of serum-supplemented DMEM without antibiotics and incubated for 12 h at 37□°C and 5% CO₂. Post incubation of the scaffolds with immune cells, bacterial cultures were washed and resuspended in PBS (pH 7.4), and a 7 µL aliquot was infused onto PVAc scaffolds immersed in the RAW cell suspension to achieve a multiplicity of infection (MOI) of 10:1 (approximately 3□×□10□ bacteria per 0.3□×□10□ macrophages). Co-incubation was performed for 24□h at 37□°C in a humidified atmosphere containing 5% CO₂. Bacterial aggregates on scaffolds were quantified by confocal microscopy, infection levels were assessed as CFUs (extracellular and internalized bacteria), and host cell responses, including bacterial phagocytosis and cell viability, were measured by flow cytometry.

For experiments involving polystyrene tubes, silicone-coated tubes (or uncoated) were UV-sterilized for 30 min, rinsed with ethanol, and washed with deionized water. Tubes were coated with 1% BSA at 37□°C for 45 min, then filled with PBS, complete medium (IMDM/RPMI + FBS), or 1 × 10□ HL-60/THP-1 cells in media and incubated at 37□°C for 8h, after which cell counts were recorded and liquids discarded. Subsequently, LB with ∼10□ CFU of *E. coli* MG1655 was added, followed by 48 h incubation. Planktonic bacteria were quantified as CFUs, and biofilm biomass was measured by crystal violet staining.

### Hyaluronic Acid – Polyarginine (HA-PA) Coating

The HA-PA coating procedure was adapted from Mutschler et al.^32^ HA and PA were prepared at a concentration of 0.5 mg/mL in a buffer containing 150 mM NaCl and 10 mM Tris (pH 7.4). Sectioned PVAc scaffolds were sequentially immersed in HA solution (200 µL) for 1 minute and PA solution (200 µL) for 1 minute in a 48 well plate. This alternating immersion cycle was repeated six times to achieve six layers of HA and PA on the PVAc scaffold, denoted as (HA-PA)₆.

### Silicone and Silicone+ Coatings

PVAc scaffolds/PS tubes were coated with food grade silicone (DOWSIL 734 flowable sealant, Dow Corning, USA) and allowed to cure at room temperature for 1 hour. Following curing, the silicone-coating was sterilized under ultraviolet light for 1 hour and then subsequently rinsed with 100% ethanol, followed by PBS. For silicone coating incorporating Bismuth Telluride (Bi_2_Te_3_), the compound was mixed with Silicone at a concentration of 15 mg/mL. For the silicone incorporating HA-PA coating, the procedure described for the HA-PA coating (above) was followed following the silicone-coating of PVAc.

### Synthesis of Bi_2_Te_3_ hexagonal nanoflakes

A precursor medium was first obtained by dispersing sodium hydroxide (0.32 g) together with polyvinylpyrrolidone (0.44 g) in ethylene glycol (24 mL) under continuous magnetic agitation at ambient conditions for 24 h. Subsequently, bismuth nitrate pentahydrate (0.38 g) and sodium telluride (0.26 g) were introduced into the prepared mixture, followed by further mixing for 5 h to achieve a uniform reaction system. The resulting suspension was then poured into a Teflon lined vessel, sealed within a stainless-steel autoclave and heated at 200 °C for 36 h in a laboratory oven. After completion of the solvothermal process, the product was collected by centrifuging at 12,000 for 10 min. The recovered solid was purified sequentially using deionized (DI) water and absolute ethanol. Each cleaning step was repeated three times under identical centrifugation conditions. Finally, the collected material was rapidly frozen in liquid nitrogen and dried in a freeze dryer for 24 h, yielding Bi_2_Te_3_ powder composed of hexagonal nanoflakes.

### Measurement of H_2_O_2_ generation from Bi_2_Te_3_

Bi_2_Te_3_ hexagonal nanoflakes were dispersed in 5 mL of DI water and sealed with aluminum foil to eliminate light exposure during the experiment. The prepared vials were subjected to thermal stimulation in a water bath under heating conditions of 27, 30 and 33 °C, followed by a cooling step at 20 °C. During a single heating/cooling sequence, each vial remained in the water bath for 6 min. The thermal cycling procedure was repeated five times. Hydrogen peroxide production was subsequently quantified after completion of the five cycles under temperature gradients (ΔT) of 7, 10, and 13 °C. Hydrogen peroxide generation was quantified using Amplex Red/horseradish peroxidase (HRP) fluorescence assay. Two precursor solutions were prepared prior to the measurements. The first solution consisted of 0.4 mg Amplex Red dissolved in 3.1 mL DMSO, while second solution was prepared by dissolving 0.5 mg HRP in PBS. For the catalytic experiments, Bi_2_Te_3_ samples were dispersed in 5 mL DI water and exposed to controlled temperature treatment in a water bath for 6 min. Following thermal stimulation, the suspension was passed through a 0.2 µm polyvinylidene fluoride (PVDF) membrane to remove solid particles. Subsequently, 270 µL of the collected filtrate was mixed with 30 µL of Amplex Red solution and 3 µL of HRP solution. Fluorescence measurements were carried out using Hitachi High-Tech F-7000 photoluminescence spectrophotometer. The reaction mixture was excited at 530 nm and the emission spectrum was recorded over a wavelength range of 560-750 nm.

### Experiments Involving Peripheral Venous Blood Draws

#### Human Ethics Statement

All research involving human samples was conducted in accordance with protocols approved by the Human Ethics Committee of the Indian Institute of Science (approval numbers IHEC No: 19-14012020 and IHEC No: 04/05042024). Informed consent was obtained from all volunteer donors prior to sample collection.

On the day of experimentation, 3 mL of peripheral venous blood was collected from healthy volunteers and transferred immediately into EDTA-coated tubes. Of this, 2 mL was carefully layered along the walls of a 15 mL Falcon tube containing 3 mL of Histopaque-1077 (Merck) at a 60° angle. The samples were centrifuged at 500g for 20 minutes at room temperature without braking, enabling density-based separation of blood components. The plasma fraction was recovered at the top, with a distinct buffy coat (PBMCs) located between the plasma and Histopaque layer, and a pellet containing PMNLs and RBCs at the bottom. The buffy coat and pellet fractions were transferred into separate 15 mL tubes.

Red blood cell lysis buffer (150 mM NH₄Cl, 10 mM NaHCO₃, 1.2 mM EDTA) was added to each tube (approximately 1 mL for PBMCs and 10 mL (1:10 ratio) for PMNLs) and incubated for 10 minutes at room temperature. Samples were centrifuged at 500g for 5 minutes at 4 °C, and the resulting cell pellets were resuspended in DMEM supplemented with 10% FBS. Cells were counted, and 1 × 10^6^ PBMCs or PMNLs in 2 mL complete media were used for immune surveillance assays. For whole blood experiments, 1 mL of blood diluted with 4% sodium citrate was utilized.

### Assessment of Extracellular and Intracellular Bacteria

To quantify planktonic extracellular *S. aureus* during infection assays, 50 µL of the co-incubation culture medium was serially diluted and plated on TSB agar supplemented with chloramphenicol. Plates were incubated at 30□°C for 24 hours to allow colony formation, reported as colony-forming units (CFU). Intracellular bacteria within RAW macrophages were determined by lysing host cells with 0.1% Triton X-100, followed by plating of the lysate.

### Bacterial Uptake and Live Cell Assessment by Flow Cytometry

Phagocytosis of *sGFP*-expressing *S. aureus* was assessed using flow cytometry by gating host cells positive in the 488/520 (excitation/emission) channel. Additionally, host cells were stained with propidium iodide (PI; Merck, USA) to stain the membrane-compromised cells. The proportion of PI-negative cells was reported as the percentage of live cells.

### Confocal Imaging

Images of PVAc scaffolds colonized with GFP-labelled *S. aureus* or *E. coli* aggregates were acquired using a Leica SP8 confocal microscope equipped with a 63x oil immersion objective, with excitation at 488 nm and emission at 515 nm. Following infection, the PVAc scaffolds were carefully transferred into 35 mm glass-bottom dishes, and 20 single-plane images with 1 µm z-slices were collected across 4-6 distinct fields of view (FOV). For *in-vivo* studies, 63 single-plane images with 1 µm z-slices were collected at 6-8 fields of view. Image processing was performed in Fiji (Image J), where single-plane images were stacked to generate Z-stack reconstructions. Quantitative analysis was conducted using a custom Fiji script to determine the average area and cumulative GFP signal of all events (individual cells or multicellular aggregates), reported as *Aggregates size* and *Sum of all aggregates*, respectively. To visualize the colocalization of the bacterial aggregates with membrane compromised host cells, the scaffolds was incubated in PI (2 µg/mL in PBS) for 10 minutes and then examined under microscope.

### Crystal Violet Assay

To quantify biofilm formation on the inner surfaces of silicone-coated (or uncoated) polystyrene tubes, crystal violet staining was employed. Briefly, 1 mL of 0.25% crystal violet solution was introduced into each tube to stain the biofilms. Uniform staining was ensured by placing the tubes on a rotor for 30 minutes. Excess dye was removed by thorough washing with deionized water, followed by air drying. Subsequently, 1 mL of 33% glacial acetic acid was added to each tube, and the tubes were rotated for 30 minutes to facilitate de-staining. The absorbance of the resulting solution was measured at 590 nm, providing a quantitative assessment of biofilm biomass on the tube surfaces.

### Measurement of Extracellular Superoxide Production by RAW 264.7 Macrophages

RAW 264.7 macrophages were seeded in 96-well plates at densities of 0.3 × 10□ or 0.9 × 10□ cells per well in 200 µL of 10% DMEM (cell-free wells served as controls). The cells were allowed to adhere overnight at 37□°C in a humidified 5% CO₂ incubator. Following this, media supernatant was removed and fresh solution (250 µL Hank’s Balanced Salt Solution) containing cytochrome c (working concentration of 75 µM), with and without superoxide dismutase (SOD; working concentration of 120 U/mL) and Bi_2_Te_3_, was added. SOD was added to account for baseline superoxide levels. Following addition of reaction mixture, absorbance was taken immediately (0 h; baseline correction) and 4 hours post incubation at 540, 550, and 560 nm. Non-specific absorbance values at 540 and 560 nm were averaged and subtracted from the specific absorbance at 550 nm to obtain normalized values. Extracellular superoxide (O₂⁻•) concentrations were subsequently calculated from the normalized absorbance.

### Scanning Electron Microscopy

PVAc scaffolds were fixed in 4% formaldehyde in PBS (pH 7.4) for 15 minutes and then air-dried. The scaffolds were subsequently mounted on aluminium stubs using double adhesive carbon tape. For polystyrene tubes, the liquids inside the tubes were discarded and the tubes were washed with deionized water. Biofilms on the surface were fixed using a 1:1 solution of 10% glutaraldehyde and 10% formaldehyde and then spun on a rotor overnight at RT. The tube was then cut into small pieces and mounted on Aluminium stubs.

All the prepared samples were desiccated for 24 hours before being sputtered with gold using a BAL TEC SCD 005 (BAL-TEC, USA) sputter coater and then examined on APREO 2S HiVac SEM (ThermoFisher, USA).

### In Vivo Experiments

#### Animal Ethics Statement

Animal experiments were performed in accordance with the Guidelines for the Control and Supervision of Experiments on Animals (CPCSEA), 1998, under the Ministry of Environment and Forests, Government of India, as well as the regulations of the Institutional Animal Ethics Committee (IAEC), IISc. The study protocol was reviewed and approved by the Committee for the Purpose of Control and Supervision of Experiments on Animals (CPCSEA) at IISc (approval number: CAF/ETHICS/168/2025).

Female C57BL/6 mice (8–10 weeks old) were procured from the Central Animal Facility at IISc and acclimatized for two weeks prior to surgical procedures. Anesthesia was induced and maintained using isoflurane (Baxter, USA) through an anesthesia-delivery machine (Orchid Scientific, India). The dorsal region of each mouse was shaved and disinfected with Povidone-Iodine followed by 70% isopropyl alcohol. A small incision was made at the sterile site, and PVAc scaffolds (approximately 4 × 4 × 3 mm³) were implanted into the dorsal subcutaneous space (two scaffolds per mouse). The incision was then closed using silk sutures (Ethicon, India), and postoperative analgesia was provided with meloxicam.

Infection was performed right after surgery (0 day), 5-days and 12-days post-surgery to model no (simultaneous), moderate, and high immune cell infiltration on the scaffolds, respectively. For infection, 25 µL of *S. aureus* suspension in PBS (pH 7.4), corresponding to 2 × 10□ CFU, was injected into each scaffold. After 24 hours, mice were euthanized by CO_2_ asphyxiation, and the scaffolds were removed, fixed in 4% formaldehyde (in PBS, pH 7.4), and examined for bacterial aggregates using confocal microscopy and scanning electron microscopy (SEM). Experiments were performed where two numbers of each biomaterial were implanted into a single mouse.

## Acknowledgements

This work was partly funded by the India-Taiwan Programme of Cooperation in Science & Technology (Indian National Academy of Engineering, Department of Science and Technology India and Department of International Cooperation and Science Education, National Science and Technology Council, Taiwan) awarded to Z-HL and SJ (reference number 2024/IN-TW/06). Additional funding for this work was through the DBT-Wellcome India Alliance Intermediate fellowship to SJ (reference number IA/I/19/1/504265). We also acknowledge Ravi Khetan for their generous contribution in memory of late Pushpa Khetan, which helped partly fund this research

## Declaration

The authors have no conflicts of interest to declare.

The authors declare that they have used the assistance of a generative AI tool (Claude, Anthropic) for editing and improving clarity of text. The tool was not used for data generation, data analysis or interpretation of results. The authors also declare that all AI-assisted text was reviewed and further edited by them and take full responsibility for the content presented here.

## Notes

### Competing Interest Statement

The authors have declared no competing interest.

### Summary of Updates

Additional data and models have been added to this revision. These were done to improve the quality of our work and to explain specific mechanistic aspects of the research. Additionally, an in vivo model was added to demonstrate the potential real-world application of our data. All of these data were generated by additional personnel, and hence the author list has changed substantially. However, the fundamental concept of the manuscript and the main conclusions remain the same.

