## Supplementary for "Immune Cell Attachment on Material Surface Promotes Bacterial Aggregation and Biofilms"

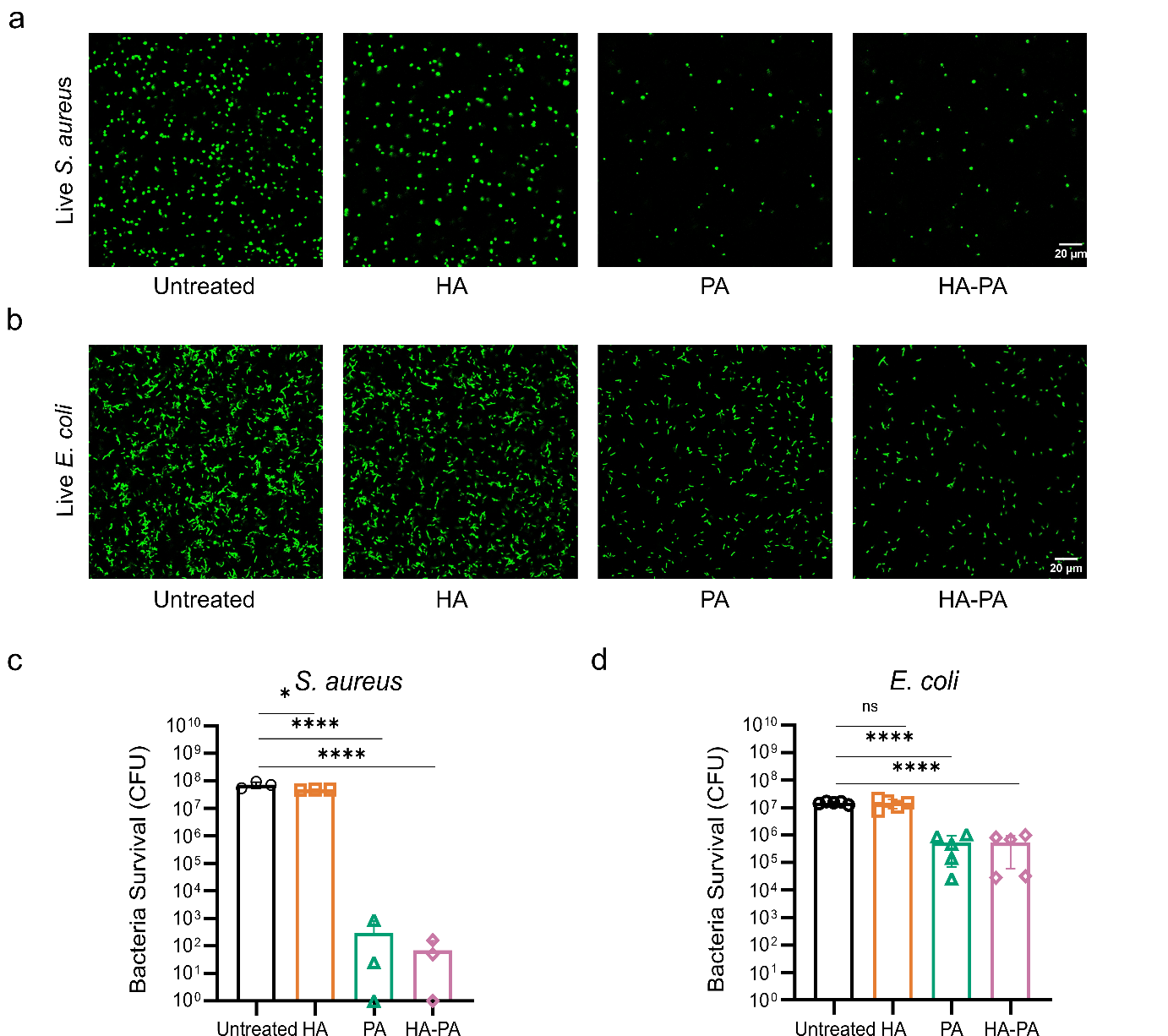

**Figure S1. Antibacterial capability of HA, PA, and HA-PA against planktonic bacterial cultures of *S. aureus* and *E. coli.***

(a) and (b) – Representative confocal images of *S. aureus* (a) and *E. coli* (b) cultures following various treatments (n=3).

(c) and (d) Quantification of live *S. aureus* (c) and *E. coli* (d) expressed as colony forming units (CFU).

Statistical analysis was performed using one-way ANOVA followed by Dunnett’s multiple comparison test. * *p<*0.05, **** *p<*0.0001, and ns=non-significant.

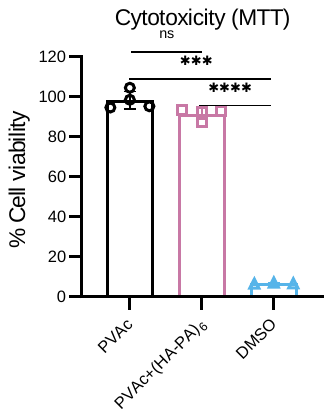

**Figure S2. Cellular cytotoxicity assessment of PVAc scaffolds with and without HA/PA coating.**

RAW 264.7 macrophage viability was measured using MTT assay (measuring absorbance at 595 nm) and values were normalized to negative control (cells without scaffolds, 100%). DMSO was used as a positive control (as it causes cell death). One way ANOVA followed by Tukey’s multiple comparison test. *** *p<*0.001, *****p<*0.0001, and ns=non-significant.

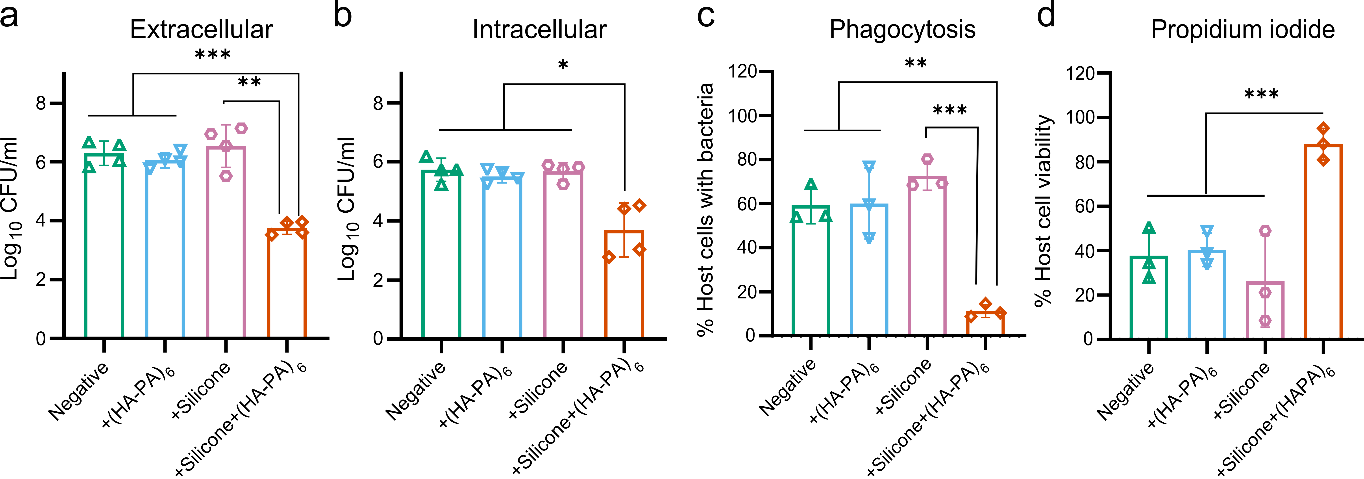

**Figure S3.** **Effect of silicone and silicone-HA-PA coating on *S. aureus* clearance from biomaterial surfaces*.***

(a) Enumeration of planktonic extracellular surviving *S. aureus* in coculture supernatant expressed as CFU.

(b) Evaluation of internalized surviving *S. aureus* recovered from macrophages in coculture expressed as CFU.

(c) Flow cytometric quantification of the percentage of live macrophages with GFP-expressing *S. aureus*.

(d) Flow cytometric quantification of the percentage of live macrophages following infection and treatment conditions.

All CFU and flow cytometry results were presented as mean ± SD (n=5 for CFU; n=3 for flow cytometry). Statistical analysis was performed using one way ANOVA followed by Tukey’s multiple comparison test. Statistical significance is indicated as **p<*0.05, ***p<*0.01, ****p<*0.001, and ns=non-significant.

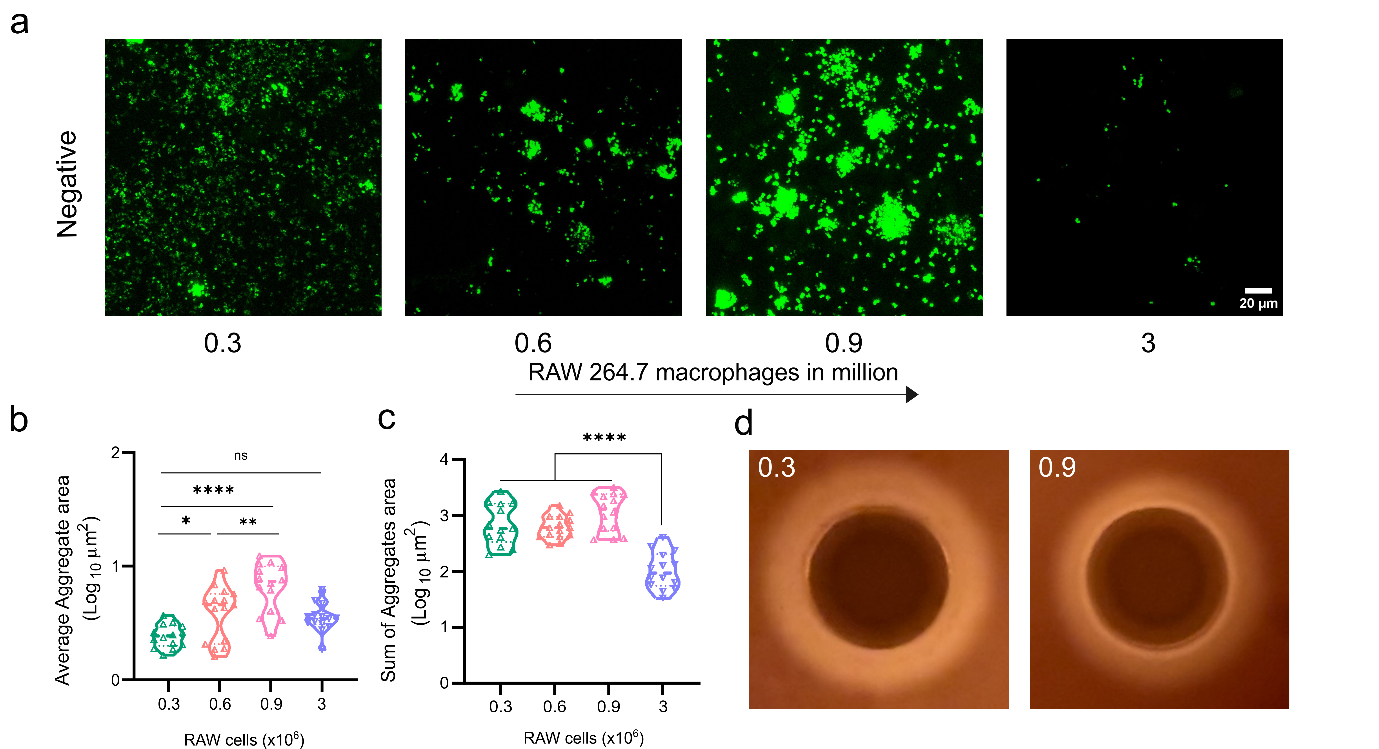

**Figure S4. Effect of enhanced macrophage interaction with biomaterial on *S. aureus* aggregation.**

(a) Representative confocal microscopy images showing *S. aureus* aggregation on biomaterial surfaces as a function of increasing macrophage numbers. Negative implies PVAc scaffold without any coating.

(b) Quantitative analysis of bacterial aggregate area, in response to varying macrophage densities.

(c) Quantification of total bacterial aggregates in response to varying macrophage densities.

(d) δ-hemolysin secretion assessed using a sheep blood agar plate demonstrates a reduced zone of clearance at higher macrophage numbers, indicating diminished *agr* expression and concomitant upregulation of biofilm-associated factors.

Bacterial aggregate quantification was performed using data from three independent experiments with minimum 4 FOVs analyzed per biomaterial. Statistical analysis was performed using one way ANOVA followed by Tukey’s multiple comparison test, and two-way Anova with Sidak’s multiple comparison test. Statistical significance is indicated as **p<*0.05, ***p<*0.01, ****p<*0.001, *****p<*0.0001, and ns=non-significant.

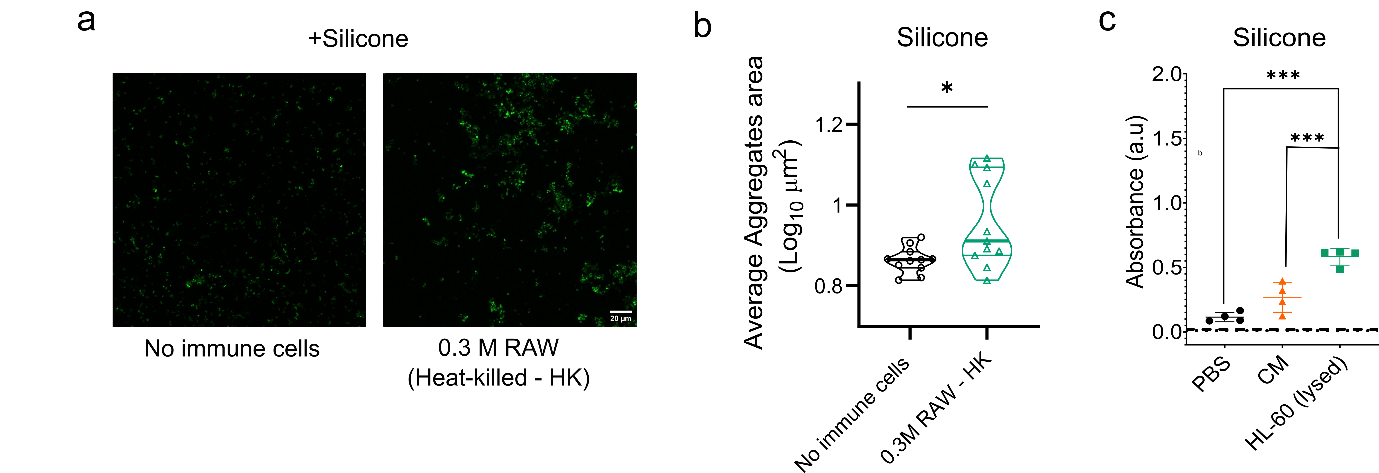

**Figure S5. Effects of RAW 264.7 macrophage on *S. aureus* aggregation and HL-60 lysates on *E. coli* biofilm formation on biomaterial surfaces.**

(a) Representative confocal microscopy images showing *S. aureus* aggregation on silicone surface in the presence of heat-killed macrophages

(b) Quantitative analysis of bacterial aggregate area on Silicone surface following exposure to heat-killed macrophages

(c) Quantitative analysis of *E. coli* biofilm using crystal violet assay after incubation with HL-60 lysates in silicone tubes

Bacterial aggregate quantification was performed using data from three independent experiments with minimum 3 FOVs analyzed per biomaterial. Statistical analysis was performed using unpaired t-test followed by Mann-Whitney test, and one-way Anova with Tukey’s multiple comparison test. Statistical significance is indicated as **p<*0.05, ***p<*0.01, ****p<*0.001, *****p<*0.0001, and ns=non-significant.

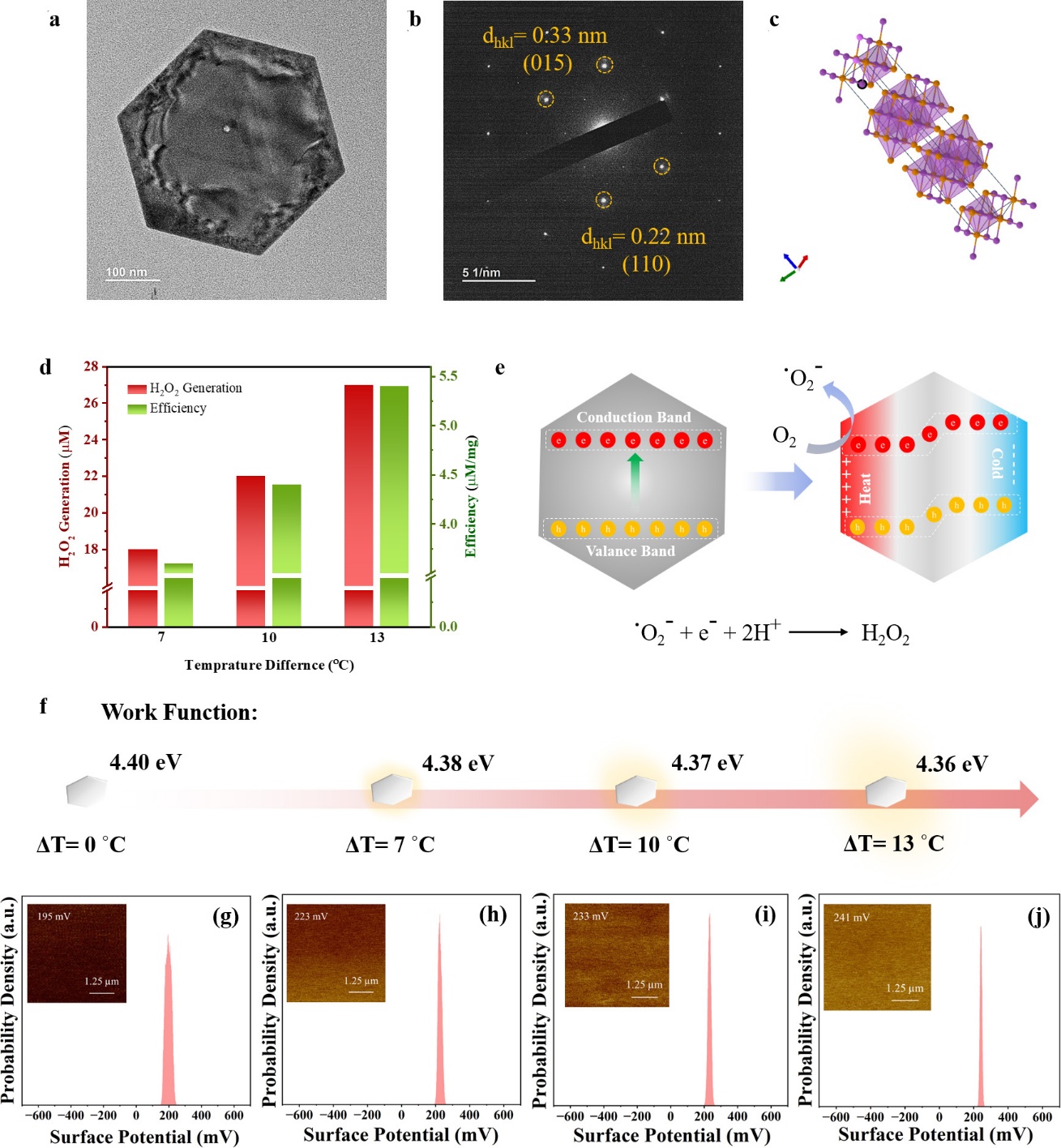

**Figure S6.** **Structural characterization, thermally induced charge modulation and temperature-gradient driven H_2_O_2_ generation of synthesized hexagonal nanostructures.**

(a) transmission electron micrograph (TEM) of the Bi_2_Te_3_ nanoflakes

(b) SAED pattern showing the morphology of the hexagonal nanosheet

(c) Crystal structure illustrating layered atomic arrangement

(d) H_2_O_2_ generation and catalytic efficiency under different temperature differences (ΔT = 7, 10 and 13 °C)

(e) Schematic illustration of the thermally induced charge separation mechanism and oxygen reduction pathway for H_2_O_2_ generation under temperature gradient

(f) Work function values obtained for Bi2Te3 at varying temperature difference

(g-j) KPFM surface potential mapping and corresponding gaussian distributions measured at different temperature differences.

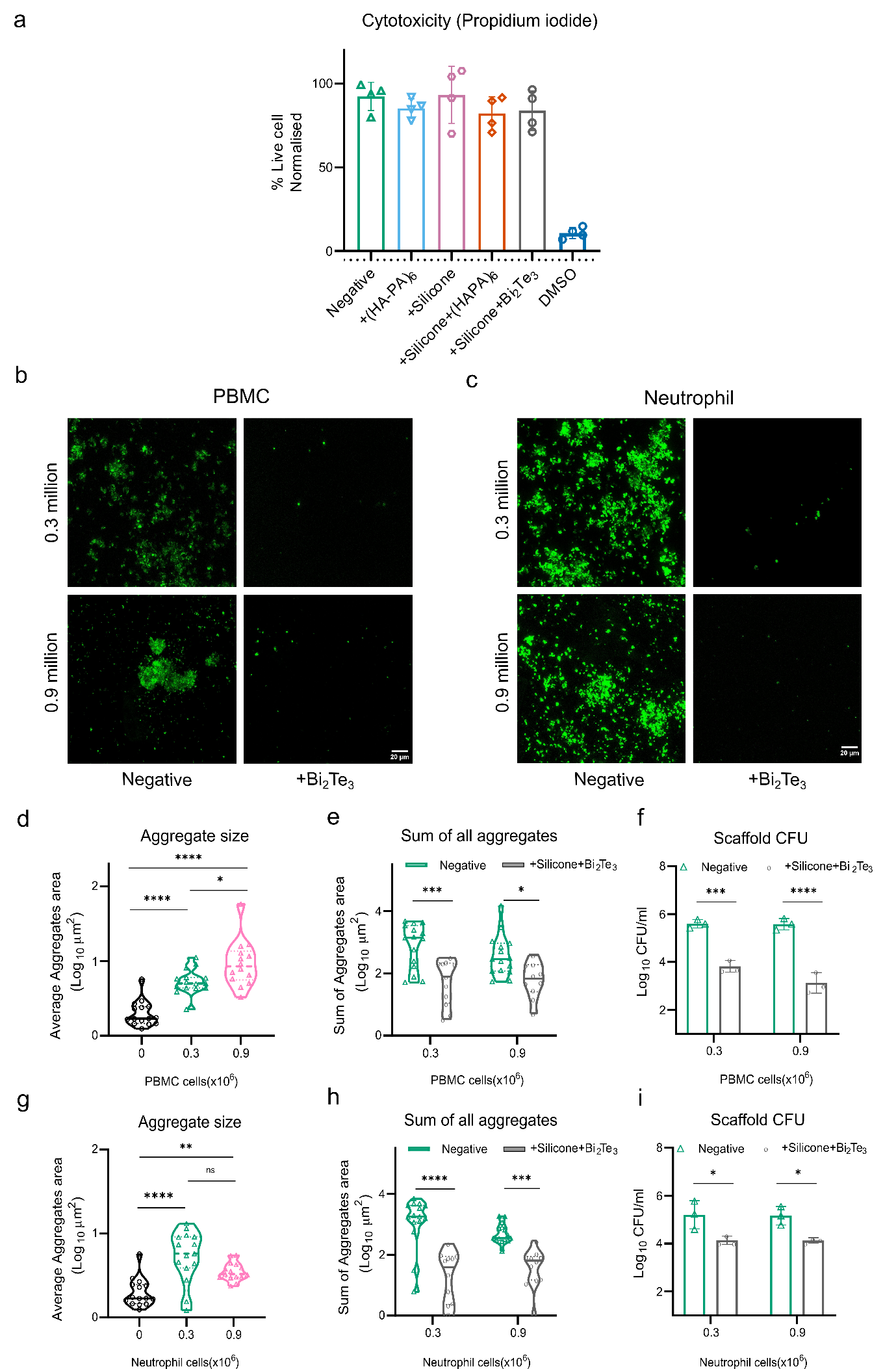
**Figure S7. Influence of human PBMCs and neutrophils on *S. aureus* aggregation and antibiofilm effect of Silicone+Bi_2_Te_3_ coating.**

(a) Cytotoxicity assessment of PVAc scaffolds with various listed surface coatings (negative implies no coating – only PVAc scaffold). Cell viability was measured using propidium iodide staining, with viable cells quantified by gating PI-negative population via flow cytometry and values were normalized with negative control (cells without scaffolds). DMSO served as positive control.

(b) and (c) Representative confocal images showing *S. aureus* bacterial aggregates in the presence of 0.3 and 0.9 million PBMCs (b) and neutrophils (c), in the absence and presence Silicone+Bi_2_Te_3_ coating.

(d) – (i): Quantification of bacterial aggregate size in the absence and presence of (d) PBMCs and (g) Neutrophils at 0.3 and 0.9 million cells; Quantification of total bacterial aggregates with and without treatment in (e) PBMCs and (h) Neutrophils; Quantification of surviving *S. aureus* as CFU recovered from the biomaterial with and without treatment in the presence of (f) PBMCs and (i) Neutrophils.

Statistical analysis was performed using one-way ANOVA followed by Tukey’s multiple comparison test and two-way Anova with Sidak’s multiple comparison test. **p<*0.05, ***p<*0.01, ****p<*0.001, *****p<*0.0001, and ns=non-significant.

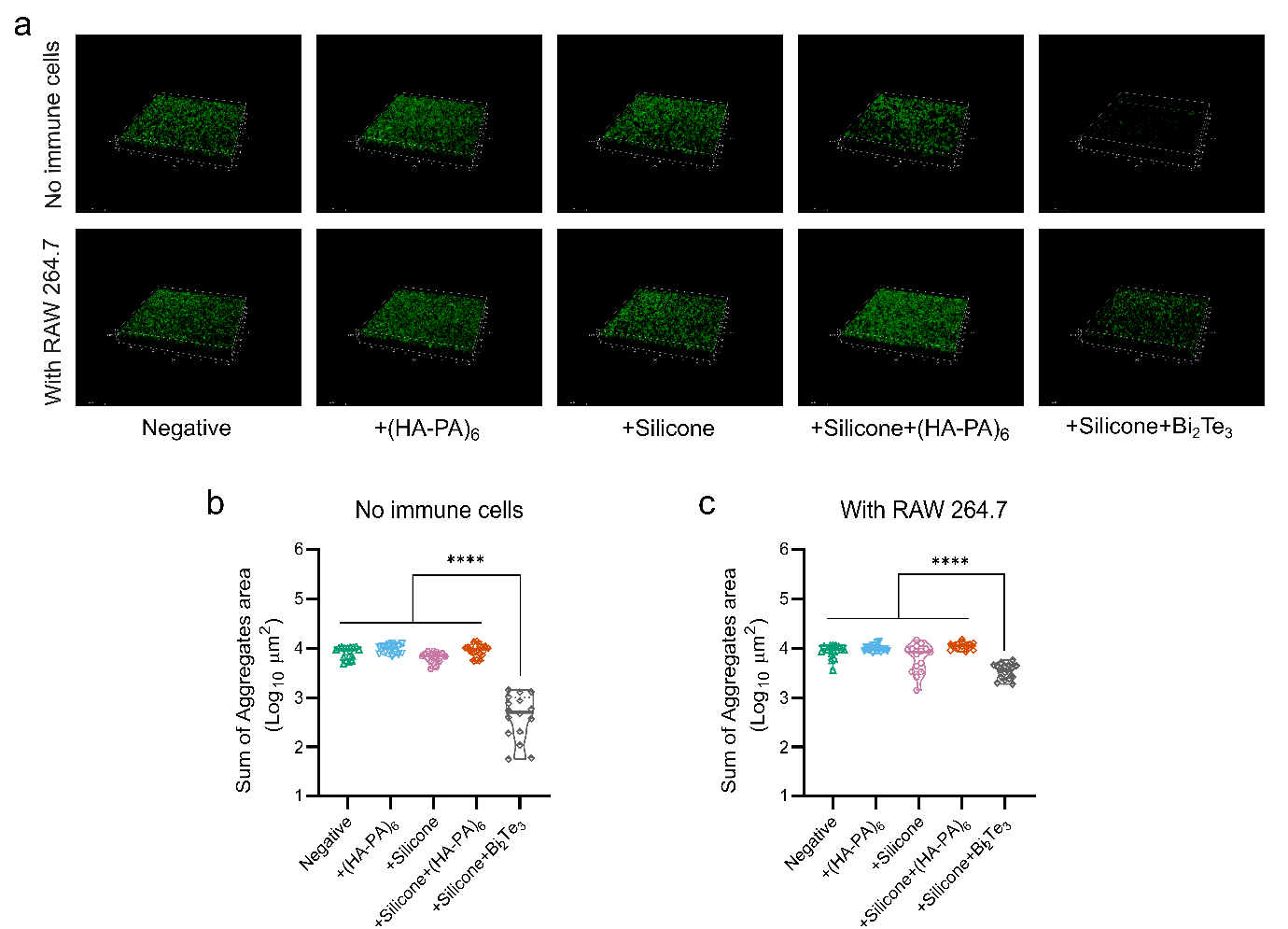

**Figure S8. Inhibition of *E. coli* biofilm formation by bismuth telluride.**

(a) Representative confocal microscopy images showing surviving *E. coli* on biomaterial surfaces under different coating conditions in absence and presence of RAW macrophages. Negative implies no coating (PVAc scaffold only)

(b) Quantitative analysis of total bacterial aggregates area in the absence of macrophages.

(c) Quantitative analysis of total bacterial aggregates area in the presence of macrophages.

Bacterial aggregate quantification was performed using data from three independent experiments with atleast 4 FOVs analyzed per biomaterial. Statistical analysis was performed using one way ANOVA followed by Tukey’s multiple comparison test. Statistical significance is indicated as *****p<*0.0001.

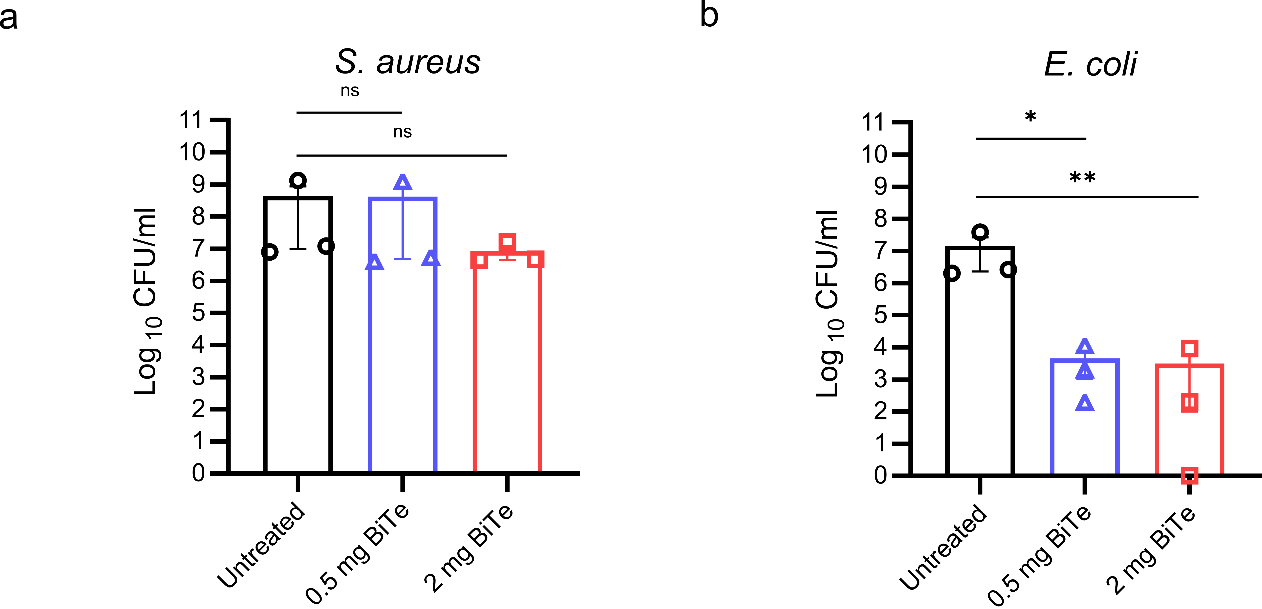

**Figure S9. Antibacterial capability of Bi_2_Te_3_ against planktonic bacterial cultures of *S. aureus* and *E. coli.***

(a) and (b) Quantification of *S. aureus* (a) and *E. coli* (b) expressed as colony forming units (CFU) at no Bi_2_Te_3_, 0.5 and 2 mg Bi_2_Te_3_ (n=3).

Statistical analysis was performed using one-way ANOVA followed by Dunnett’s multiple comparison test. **p<*0.05, ***p<*0.01, and ns=non-significant.

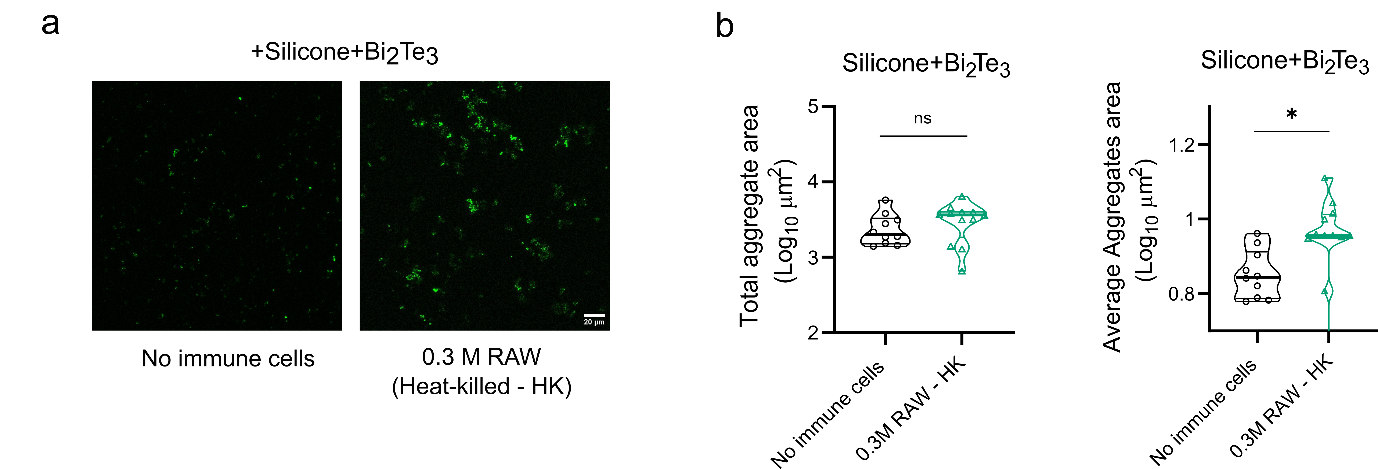

**Figure S10. Effects of RAW 264.7 macrophage lysates on *S. aureus* aggregation on Silicone+Bi_2_Te_3_ coated surfaces.**

(a) Representative confocal microscopy images showing *S. aureus* aggregation on silicone+Bi_2_Te_3_ surface in the presence of heat-killed macrophages (or no immune cells).

(b) Quantitative analysis of bacterial aggregate area (total and average) on silicone+Bi_2_Te_3_ surface following exposure to heat-killed macrophages.

Bacterial aggregate quantification was performed using data from three independent experiments with minimum 3 FOVs analyzed per biomaterial. Statistical analysis was performed using Mann-Whitney test. Statistical significance is indicated as **p<*0.05, ***p<*0.01, ****p<*0.001, *****p<*0.0001, and ns=non-significant.
